# Harnessing the *PRDM1–PGC1Α* axis to enhance CAR T cell efficacy

**DOI:** 10.64898/2026.10.15.759372

**Authors:** Sidney YY Wang, Xiufen Chen, Xubin Li, Ali ElSheikh, Alan Cooper, Joanna Chorazeczewski, Lishi Xie, Jerry YC Hu, Han Liu, Michael R Green, Justin Kline

## Abstract

Despite its promise in treating B cell malignancies, chimeric antigen receptor (CAR) T cell therapy remains limited by resistance mechanisms that impair CAR T cell stemness, persistence, and effector function. Here, we identified *PRDM1*, which encodes the transcriptional repressor, BLIMP-1, as a crucial regulator of CAR T cell terminal differentiation and exhaustion. *PRDM1* was highly enriched in exhausted-like tumor-infiltrating CAR T cells from lymphoma patients and in pre-infusion products from clinical non-responders. These findings link increased *PRDM1* expression to CAR T cell dysfunction and poor clinical outcomes. Genetic deletion of *PRDM1* significantly promoted CAR T cell proliferative capacity, memory differentiation and antitumor efficacy in both hematologic and solid tumor models. Mechanistically, *PRDM1* deletion increased the activity of PGC1α, a transcriptional coactivator central to mitochondrial biogenesis and antioxidant capacity, thereby improving mitochondrial function, oxidative stress resistance, and metabolic reserve. Depletion of PGC1α attenuated the improved efficacy of *PRDM1*-deficient CAR T cells, indicating an essential role for this pathway downstream of *PRDM1* loss. Pharmacologic activation of the PPAR–PGC1α program with a clinically approved agonist, bezafibrate, enhanced CAR T cell antitumor activity without direct tumoricidal effects. Notably, transient exposure to bezafibrate during CAR T cell manufacturing was sufficient to confer durable treatment benefits. Together, our study defines a *PRDM1*–*PGC1Α* axis that controls CAR T cell differentiation, mitochondrial fitness and therapeutic response, which provides a mechanistic and translational framework for advancing CAR T cell therapy across distinct tumor contexts.

## INTRODUCTION

CAR T cell therapy has transformed the treatment of hematologic malignancies, yet a substantial fraction of patients experience disease progression or relapse.^1–3^ Resistance to CAR T cell therapy can arise through multifaceted mechanisms, including host-related determinants, tumor cell-intrinsic characteristics, immune microenvironment, and CAR T cell fitness.^4–7^ Among these factors, rapid terminal differentiation and acquisition of dysfunctional states by CAR T cells have emerged as major challenges,^8–10^ which can occur during *ex vivo* manufacturing through activation or tonic CAR signaling,^11–13^ and continue following infusion through persistent antigen stimulation in an immunosuppressive tumor microenvironment.^4–7^

As T cells differentiate, they progressively lose self-renewal capacity, proliferative potential, and long-term persistence. Naïve/stem cell-like and central memory T cells retain greater developmental plasticity, whereas terminal effector cells exert immediate cytotoxicity but are often short lived.^14–16^ Under prolonged antigen exposure, T cells adopt an exhaustion program, characterized by reduced expansion, increased expression of co-inhibitory receptors, altered metabolism, and impaired effector functions.^17–19^ Clinically, the differentiation state of CAR T cells is increasingly recognized as a predictor of treatment outcomes. ^8–10^ For instance, infusion products enriched for memory-like features are associated with superior peak expansion, persistence, and clinical responses, while CAR T cells from non-responders commonly induce late differentiation, exhaustion, and apoptosis programs. Collectively, these findings highlight the need for strategies that prevent or delay differentiation toward short-lived effector or exhausted fates to improve therapeutic efficacy.

T cell differentiation and exhaustion are governed by dynamic transcriptional circuits that integrate antigen stimulation, co-stimulatory and inhibitory signaling, cytokine input, and metabolic cues.^16,18–20^ Several transcription factors, including TOX and NR4A family members, have been identified as essential regulators of T cell dysfunction. However, efforts to manipulate these factors have presented important complexities. TOX is central to establishment of exhausted T cell lineage but also supports formation of progenitor-like cells and persistence under chronic stimulation.^21–24^ Likewise, although NR4A-family deficiency can restore effector activity and resistance to hypofunction, its impact on memory differentiation is context dependent and varies between single versus pan-family targeting.^25–27^ Hence, an important goal is to identify regulators whose perturbation prevents or delays T cell exhaustion, while simultaneously preserving stemness for sustained immunity.

In this regard, *PRDM1*, encoding the transcriptional repressor, BLIMP-1, is well-positioned to address this need. BLIMP-1 functions at a distinct regulatory node under both acute and chronic stimulation, coupling memory suppression to terminal T cell differentiation and dysfunction.^28–31^ We therefore hypothesized that disrupting *PRDM1* would preserve memory-like fitness and support durable antitumor activity of CAR T cells. In support of this hypothesis, we observed that *PRDM1* was enriched in exhaustion-skewed CAR T cells from lymphoma biopsies of CAR T cell-treated patients. Increased *PRDM1* expression was also detected in pre-infusion CAR T cells from clinical non-responders, where memory and proliferation programs were concomitantly downregulated. These findings suggest that *PRDM1* may shape CAR T cell fate across the therapeutic trajectory during manufacturing and after infusion, contributing to poor clinical responses. Functional studies next demonstrated that *PRDM1* deletion enhanced CAR T cell expansion, early memory differentiation, effector cytokine production, and *in vivo* efficacy in hematologic and solid tumor models. Mechanistically, we identified PGC1α, a transcriptional coactivator that promotes mitochondrial biogenesis and redox homeostasis, as a key downstream effector in *PRDM1*-deficient CAR T cells. Leveraging this insight, we developed a novel pharmacologic strategy to activate the PGC1α program and improve CAR T cell performance, which, when compared to genetic modification, offers a safer and more scalable approach for clinical translation.

## RESULTS

*PRDM1* is enriched in CAR T cells exhibiting exhaustion phenotypes in relapsed/refractory large B cell lymphoma (r/r LBCL) specimens and infusion products from clinical non-responders. To examine the relationship between *PRDM1* expression and T cell differentiation in the lymphoma microenvironment, we analyzed single-cell RNA sequencing (scRNA-seq) profiles of tumors from 217 patients.^32^ We focused on CD8^+^ T cells because of their central role in antitumor cytotoxicity and susceptibility to exhaustion. Unsupervised clustering identified five major subsets characterized by distinct effector, memory, and exhaustion-associated transcriptional programs (**Figures 1A–C**). *PRDM1* was preferentially expressed in populations with the highest exhaustion scores (**Figure 1D**). Consistent with this distribution, inferred *PRDM1* regulatory activity was elevated in exhausted subsets and lowest in naïve T cells, with agreement between RNA-derived gene-based and ATAC-derived region-based analyses (**Figures 1E and 1F**). We next examined whether this correlation extended to CAR T cells after infusion by analyzing scRNA-seq data from tumor biopsies of r/r LBCL patients who were either CAR T-cell therapy–naïve or treated with axicabtagene ciloleucel (axi-cel).^33^ CAR-expressing clusters with pronounced exhaustion features showed increased *PRDM1* expression, alongside reduced memory-associated programs (**Figures 1G–1J**). These findings link *PRDM1* to exhausted states in both tumor-infiltrating CD8^+^ T cells and post-infusion CAR T cells. We next analyzed an independent LBCL cohort to assess whether *PRDM1* expression in pre-infusion CAR T cells was correlated with clinical responses.^10^ CD8^+^ CAR T cells from clinical non-responders (NR) showed significantly increased *PRDM1* transcript levels (**Figure 1K, left**) and reduced expression of *PRDM1* regulon genes (**Figure 1K, right**), indicative of *PRDM1*-mediated transcriptional repression. We further stratified patients based on *PRDM1* levels in total CAR^+^ T cells. The *PRDM1*-high group showed enrichment of exhaustion-associated pathways, accompanied by decreased memory and proliferation potential. Moreover, the MYC target program, a canonical downstream axis suppressed by *PRDM1*,^34,35^ was significantly downregulated, which supported the mechanistic impact of higher *PRDM1* expression (**Figure 1L**). Together, these findings suggest that *PRDM1* might not only regulate CAR T cell differentiation after infusion but also imprint cell fate early during manufacturing, thereby predisposing CAR T cells to reduced durability and potency upon lymphoma cell encounter, and culminating in inferior clinical outcome. We therefore examined whether disrupting *PRDM1* would improve intrinsic CAR T cell fitness and enhance antitumor efficacy.

**Figure 1.**
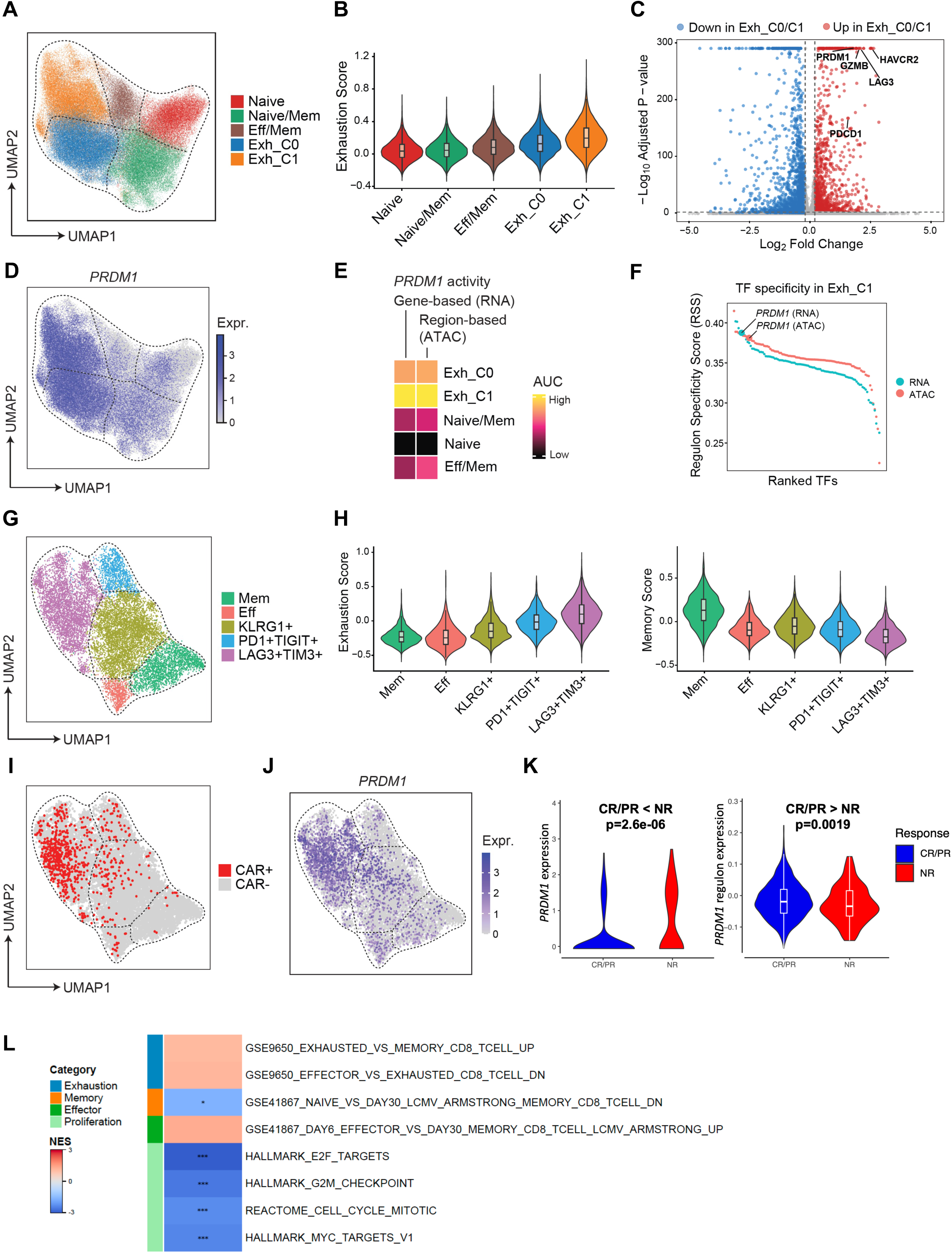
*PRDM1* is highly enriched in exhausted-like CD8^+^ CAR T cells from r/r LBCL patients and infusion products from non-responders with decreased proliferative and memory potential. **(A)** Uniform manifold approximation and projection (UMAP) plot of CD8^+^ T cells showing naive, memory (Mem), effector (Eff), and exhausted (Exh) subsets from patients with DLBCL (n=217, EGAS50000001022). **(B)** T cell exhaustion signature scores across the identified CD8^+^ T cell subsets. **(C)** Differentially expressed genes (DEGs) compared between exhausted CD8^+^ T cells and all remaining CD8^+^ T cell subsets. Dashed lines indicate statistical significance and fold-change thresholds. **(D)** *PRDM1* expression across CD8^+^ T cells. **(E)** *PRDM1* enhancer regulon (eRegulon) activity scores across indicated CD8^+^ T cell subsets. **(F)** Ranked Regulon Specificity Score (RSS) plot of candidate transcription factors (TFs) in the Exh_C1 CD8^+^ T cell subset. **(G)** UMAP plot of CD8^+^ T cell subsets from r/r LBCL tumors across CAR T naïve patients (CAR-naïve, n = 5) and those refractory to CAR T cell therapy (post-CAR, n = 7) (EGAS50000001982). **(H)** T cell exhaustion signature scores (left) and memory signature scores (right) across the identified CD8^+^ T cell clusters. **(I)** CAR expression in the CD8^+^ T cells. **(J)** *PRDM1* expression across CD8^+^ T cells. **(K)** *PRDM1* expression in CD8^+^ T cells from infusion products of patients with LBCL (GSE197268). Patient outcomes are defined by PET/CT at day 90 post CAR T cell therapy. CR, complete response; PR, partial response; NR, no response. **(L)** Gene set enrichment analysis (GSEA) comparing *PRDM1*-high (n = 3) versus *PRDM1*-low (n = 5) CAR⁺ T cells from infusion products of patients with LBCL for gene sets associated with T-cell differentiation and proliferation (GSE197268). Samples were stratified by *PRDM1* expression using a 0.35 quantile cutoff. Positive and negative normalized enrichment scores (NES) indicate enrichment in *PRDM1*-high and *PRDM1*-low cells, respectively. *P* values were adjusted using the Benjamini–Hochberg method: \**P_adj* < 0.05, \*\**P_adj* < 0.01, \*\*\**P_adj* < 0.001. Statistics for other data was based on Mann– Whitney test.

### Genetic deletion of *PRDM1* promotes proliferation, early memory differentiation and effector functions of CAR T cells

To investigate the impact of *PRDM1* deletion on CAR T cell functions, *PRDM1* was inactivated in human CD19-targeted CAR T cells using CRISPR/Cas9 (**Figure S1G**). Given the unexplored role of *PRDM1* across clinically relevant CAR constructs, we generated *PRDM1*-deficient CAR T cells in both second-generation 4-1BB (BB)- and CD28-based platforms. Next, we established an *in vitro* restimulation assay to model progressive CAR T cell dysfunction driven by persistent antigen exposure (**Figure 2A**). This system captured phenotypic features of exhausted CAR T cells, as evidenced by diminished expansion (**Figure S1A**), sustained expression of co-inhibitory receptors (**Figures S1B-S1D**), decreased cytokine production (**Figure S1E**) and impaired tumor-cell killing (**Figure S2F**). We then examined whether *PRDM1* loss improved CAR T cell activity under serial lymphoma cell challenge. *PRDM1* deletion markedly enhanced the proliferative potential of CAR T cells (**Figure 2B**), in parallel with substantial enrichment of stem cell memory/naïve-like (Tscm/naïve-like) and central memory (Tcm) T cell subsets (**Figures 2C and 2D**). *PRDM1*-deficient CAR T cells also showed increased production of multiple effector cytokines both at baseline (D0) and at the end of chronic stimulation (**Figure 2E**), suggesting that disrupting *PRDM1* enhances the intrinsic capacity of CAR T cells to mount effector responses.

**Figure 2.**
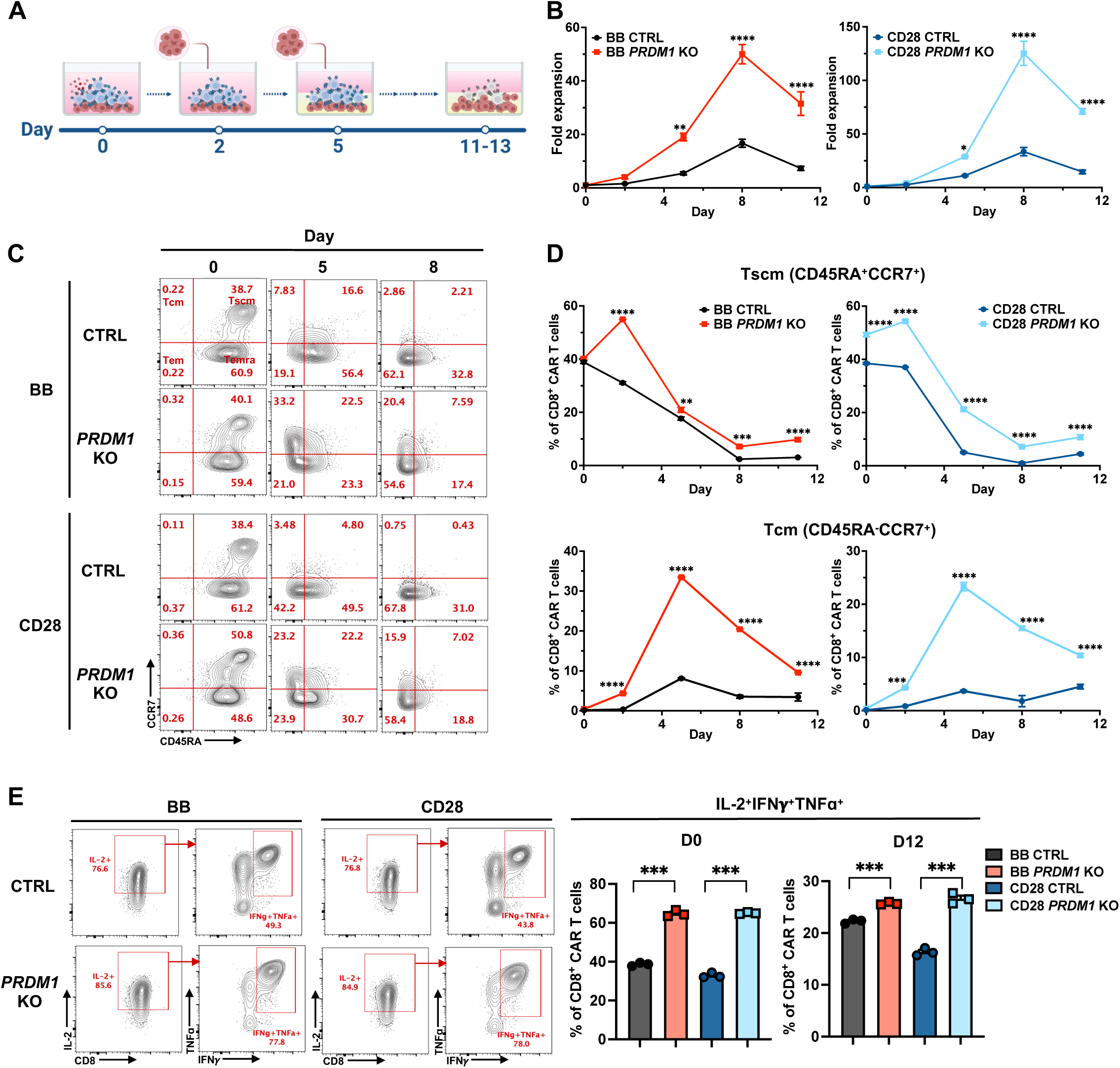
*PRDM1* deletion promotes expansion, early memory differentiation and effector function of CAR T cells. **(A)** Experimental design for *in vitro* CAR T cell restimulation assay. Control (CTRL) and *PRDM1* knock-out (KO) anti-CD19 CAR T cells containing either a 4-1BB (BB CTRL and BB *PRDM1* KO) or CD28 (CD28 CTRL and CD28 *PRDM1* KO) costimulatory domain were serially challenged with human DLBCL cell line OCI-Ly8 every 2–3 days for a total 4–5 rounds of stimulation. **(B)** Expansion kinetics of CTRL and *PRDM1* KO CAR T cells during restimulation. **(C)** Representative flow cytometry plots showing the distribution of CD8⁺ CAR T cell subsets defined by CD45RA and CCR7 expression at baseline (D0) and during restimulation. Tscm (CD45RA^+^CCR7^+^), stem cell-like memory T cells; Tcm (CD45RA^-^CCR7^+^), central memory T cells; Tem (CD45RA^-^CCR7^-^), effector memory T cells; Temra (CD45RA^+^CCR7^-^), effector memory T cells positive for CD45RA (short-lived terminally-differentiated T cells). **(D)** Frequencies of Tscm (top) and Tcm (bottom) subsets in CD8^+^ CAR T cells. **(E)** Representative flow cytometry plots (left) and frequencies (right) of CD8^+^ CAR T cells producing effector cytokines, IL-2, IFN**γ** and TNFɑ. Before and at the end of coculture, CAR T cells were restimulated with PMA/ionomycin for 5 hours in the presence of brefeldin A/monensin before analysis. Data are representative of at least 3 independent experiments using CAR T cells manufactured from different healthy donors, with 3–4 technical replicates per experiment. \**P* < 0.05, \*\**P* < 0.01, \*\*\**P* < 0.001, \*\*\*\**P* < 0.0001 using two-way ANOVA with Šídák correction for multiple comparisons (B and D) and two-tailed Welch’s *t* test (E). Error bars indicate the s.e.m.

### *PRDM1* deletion significantly improves antitumor efficacy of CAR T cells against hematologic and solid tumors

We next assessed the extent to which the advantageous *in vitro* phenotypes observed in *PRDM1*-deficient CAR T cells would translate *in vivo*. In mice xenografted with human OCI-Ly8 DLBCL cells (**Figure 3A**), *PRDM1* knock out (KO) CAR T cells demonstrated superior *in vivo* expansion (**Figures 3B and 3C**) associated with enhanced early memory subsets (**Figure 3D**). *PRDM1* deletion significantly prolonged the survival of tumor-bearing mice across both BB and CD28 costimulatory domains (**Figures 3E, 3F, S2A and S2B**). We next extended our studies to solid tumors, which remain an area of high unmet need for CAR T cell therapy. To this end, we generated HER2-directed CAR T cells against HER2-expressing human breast tumors (**Figures 4A and S3A**). *PRDM1* deletion not only increased CAR T cell expansion detected in peripheral blood (**Figures 4B and 4C**) but also infiltration into the tumor microenvironment (**Figures 4D and 4E**). Despite prolonged exposure to tumor antigen, *PRDM1* deletion maintained a higher number of early memory CAR T cells (**Figure 4F**) and preserved effector cytokine production (**Figure 4G**). Consistent with these findings, *PRDM1* disruption improved overall antitumor activity of HER2-directed CAR T cells, leading to earlier and more durable tumor control when compared to control (CTRL) CAR T cells (**Figures 4H, 4I, S3B and S3C**).

**Figure 3.**
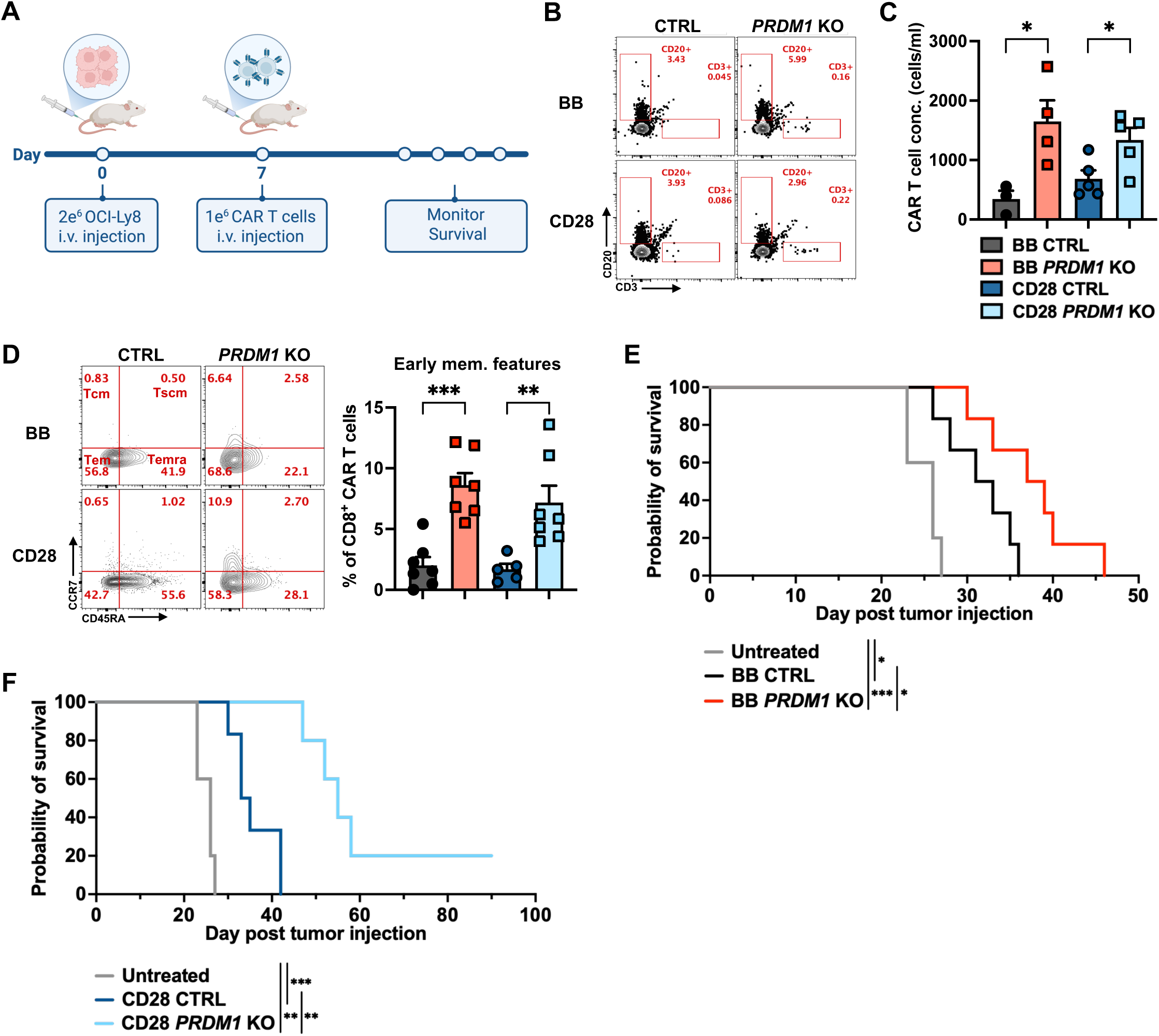
*PRDM1* deletion enhances *in vivo* expansion, memory differentiation and efficacy of CAR T cells against hematologic tumors. **(A)** Experimental design of the OCI-Ly8 xenograft mouse model. NSG mice were intravenously injected with 2 × 10^6^ OCI-Ly8 cells, followed 1 week later by infusion of 1 × 10^6^ CTRL or *PRDM1* KO anti-CD19 CAR T cells. Mice that did not receive CAR T cells served as untreated controls. **(B)** Representative flow cytometry plots showing the frequencies of human T cells in peripheral blood on D7 post CAR T cell infusion. **(C)** CAR T cell concentration in peripheral blood on D7 post infusion (n = 3–5 per group). (D) Representative flow cytometry plots (left) and frequencies (right) of CD8^+^ CAR T cells with early memory phenotypes (combined Tscm and Tcm subsets) in spleens harvested at the study endpoint (n = 5–7 per group, pooled from 2 independent experiments). **(E and F)** Kaplan-Meier survival curves following OCI-Ly8 transplantation (n = 5–6 per group). Data are representative of at least 2 independent experiments (C, E and F) using CAR T cells manufactured from different healthy donors unless otherwise indicated. \**P* < 0.05, \*\**P* < 0.01, \*\*\**P* < 0.001, \*\*\*\**P* < 0.0001 using two-tailed unpaired *t* test (C and D) and the log-rank (Mantel–Cox) test for survival analysis (E and F). Error bars indicate the s.e.m.

**Figure 4.**
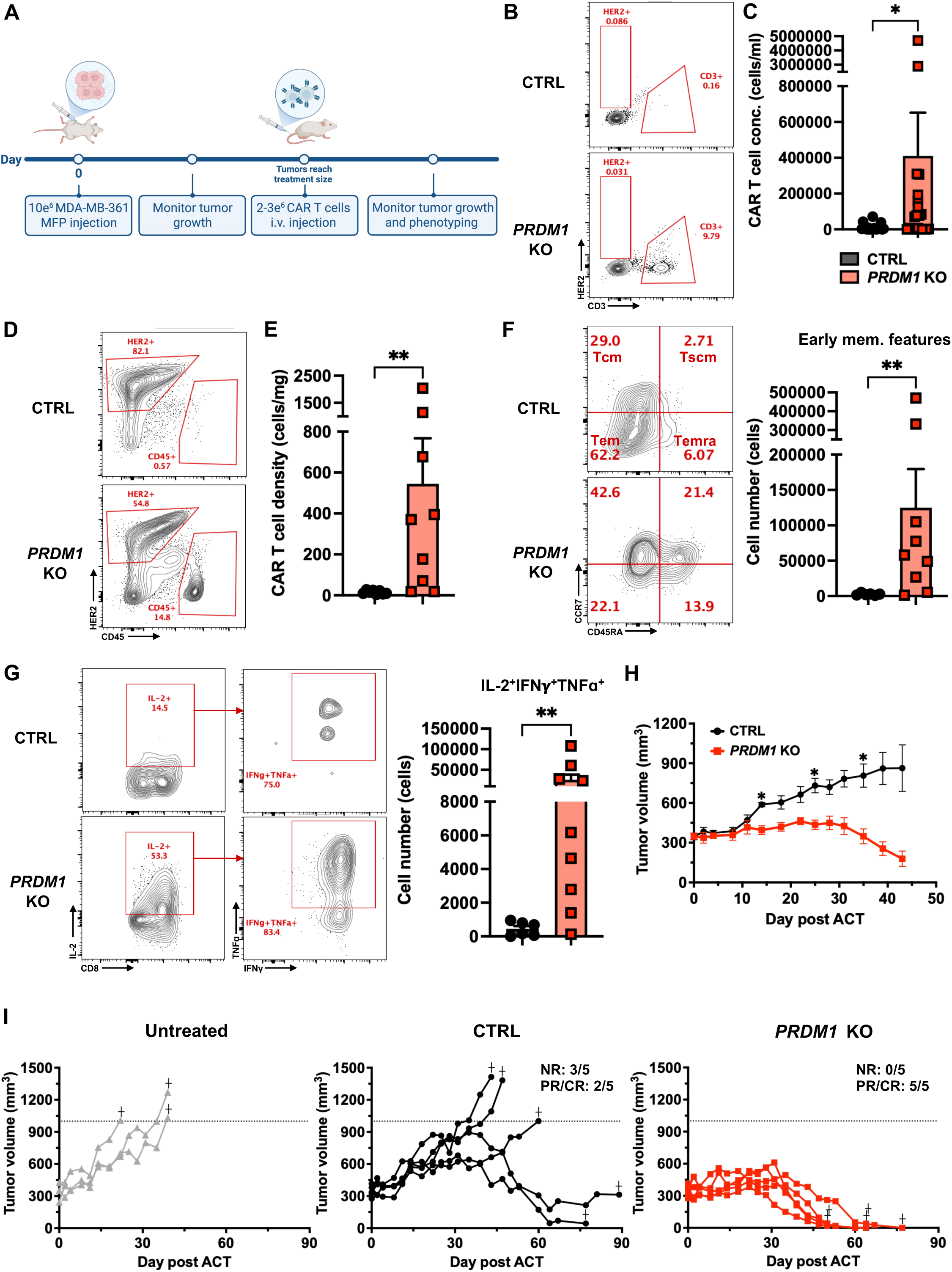
*PRDM1* deletion promotes CAR T cell infiltration, early memory differentiation, and functional persistence in solid tumors. **(A)** Experimental design of the human breast cancer cell line MDA-MB-361 xenograft mouse model. Female NSG mice received mammary fat pad injection of 1 × 10^7^ MDA-MB-361 cells. When the average tumor volume reached 200–350 mm^3^, mice were treated with 2 to 3 × 10^6^ CTRL or *PRDM1* KO anti-HER2 CAR T cells intravenously. Mice that did not receive CAR T cells served as untreated controls. Mice were euthanized when tumors exceeded 1500 mm^3^ or 1000 mm^3^ with ulceration or bleeding. **(B)** Representative flow cytometry plots showing the frequencies of human T cells in peripheral blood 5 weeks post CAR T cell infusion. **(C)** CAR T cell concentration in peripheral blood 4–5 weeks post infusion (n = 19–22 per group, pooled from 3 independent experiments). **(D)** Representative flow cytometry plots showing the frequencies of human CD45^+^ cells derived from a tumor harvested 4–5 weeks post CAR T cell infusion, corresponding to the transferred T cell population. **(E)** CAR T cell density in tumors harvested 4–5 weeks post infusion (n = 6–9 per group, pooled from 2 independent experiments). **(F)** Representative flow cytometry plots (left) and absolute number (right) of CD8^+^ CAR T cells with early memory phenotypes (combined Tscm and Tcm subsets) in tumors harvested 4–5 weeks post infusion (n = 6–9 per group, pooled from 2 independent experiments). **(G)** Representative flow cytometry plots (left) and absolute number (right) of CD8^+^ CAR T cells producing IL-2, IFN**γ** and TNFɑ. Cells were obtained from tumors harvested 4–5 weeks after infusion and restimulated with PMA/ionomycin for 5 hours in the presence of brefeldin A/monensin before analysis (n = 6–9 per group, pooled from 2 independent experiments). **(H and I)** Mean (H) and individual (I) tumor growth curves following adoptive cell transfer (ACT). The dashed line indicates a tumor volume of 1,000 mm³, above which mice were evaluated for euthanasia according to the criteria described in (A). NR, no response; PR/CR, partial or complete response. Data are representative of 2 independent experiments (n = 3–5 mice per group). Data points show measurements from individual mice. \**P* < 0.05, \*\**P* < 0.01, \*\*\**P* < 0.001, \*\*\*\**P* < 0.0001 using two-tailed Mann–Whitney test (C, E, F and G) and two-way repeated-measures ANOVA with Geisser–Greenhouse and Šídák correction (H). Error bars indicate the s.e.m.

### *PRDM1* KO CAR T cells exhibit enhanced mitochondrial, metabolic, and antioxidant potential

To gain mechanistic insight into the enhanced proliferative and functional capacity of *PRDM1* KO CAR T cells, we performed bulk RNA-seq on CAR T cells harvested during repeated stimulation. Transcriptomic analysis revealed coordinated enrichment of mitochondrial, metabolic, and antioxidant programs in *PRDM1* KO CAR T cells (**Figure 5A**). Supporting this observation, reanalysis of independent datasets from prior *PRDM1* loss-of-function CAR T cell studies recapitulated a similar transcriptional shift (**Figure S4A**). Moving forward, we examined whether the transcriptional programs enriched in *PRDM1* KO CAR T cells conferred functional mitochondrial and metabolic advantages. To assess mitochondrial resilience, CTRL and *PRDM1* KO T cells were exposed to antimycin A, which induces mitochondrial stress by inhibiting cellular respiration. CTRL T cells showed a dose-dependent increase of mitochondrial superoxide stained by MitoSOX, whereas *PRDM1* KO T cells maintained lower MitoSOX levels at baseline and after antimycin A treatment (**Figure 5B**). We then evaluated antioxidant capacity during *in vitro* restimulation, a setting in which redox imbalance typically arises and contributes to T cell exhaustion. Here again, while CTRL T cells exhibited sustained accumulation of mitochondrial superoxide, *PRDM1* KO T cells effectively attenuated mitochondrial stress throughout the experiment (**Figure 5C**). MitoTracker Green and MitoTracker Deep Red staining further showed that *PRDM1* deletion significantly increased CAR T cell mitochondrial mass and potential (**Figure 5D**). Ultrastructural imaging also corroborated a higher mitochondrial density in *PRDM1* KO CAR T cells despite repeated tumor cell challenge (**Figures 5E and 5F**). We next performed Seahorse analysis to examine mitochondrial bioenergetic capacity. Notably, *PRDM1* loss enhanced spare respiratory capacity and energetic state of CAR T cells (**Figures 5G and 5H**). Collectively, these findings indicate that *PRDM1* KO CAR T cells are equipped with improved mitochondrial fitness and metabolic plasticity to address stress and satisfy the high energy demands required during rapid expansion and eradication of target cells.

**Figure 5.**
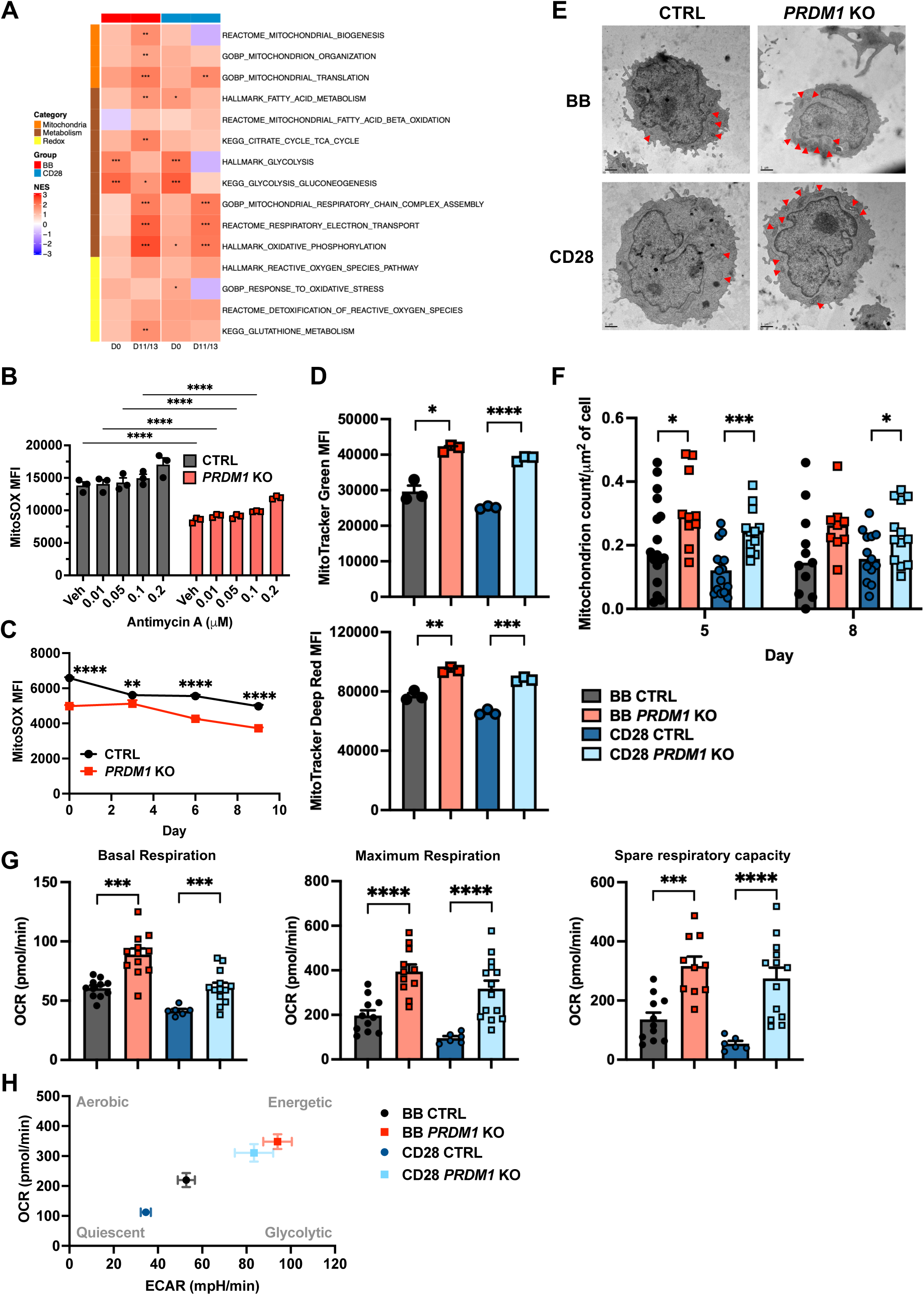
*PRDM1* deletion enhances mitochondrial, metabolic, and antioxidant potential in CAR T cells. **(A)** GSEA of *PRDM1* KO versus CTRL CD8^+^ CAR T cells containing a BB or CD28 costimulatory domain for gene sets associated with mitochondrial, metabolic, and antioxidant activity at the indicated time points. CD8^+^ CAR T cells manufactured from 3 healthy donors were sorted before and at the end of coculture with OCI-Ly8 cells for bulk RNA-seq. NES, normalized enrichment score. *P* values were adjusted using the Benjamini–Hochberg method: \**P_adj* < 0.05, \*\**P_adj* < 0.01, \*\*\**P_adj* < 0.001. **(B)** Geometric mean fluorescence intensity (gMFI) of MitoSOX in CTRL and *PRDM1* KO T cells challenged with antimycin A for 3 days. Vehicle control (veh) was treated with ethanol. **(C)** MitoSOX gMFI in CTRL and *PRDM1* KO T cells during repeated stimulation with plate-bound anti-CD3 antibodies. **(D)** MitoTracker Green and Deep Red gMFI in CD8^+^ CAR T cells repeatedly stimulated by OCI-Ly8 cells over a week. **(E)** Representative electron microscope images of CAR T cells after 5 days of repeated stimulation with OCI-Ly8 cells. Red arrows indicate mitochondria. Scale bar, 1 μm. **(F)** Quantification of mitochondrial density in CAR T cells after 5 and 8 days of repeated stimulation with OCI-Ly8 cells. **(G)** Basal respiration, maximum respiration, and spare respiratory capacity of repeatedly stimulated CD8^+^ CAR T cells measured using the Seahorse Cell Mito Stress Test. **(H)** Metabolic states (energetic, glycolytic, aerobic and quiescent) of repeatedly stimulated CD8^+^ CAR T cells assessed using the Seahorse Mito Stress Test. Data are representative of 2 independent experiments (B–H) with at least 3 technical replicates using CAR T cells manufactured from different healthy donors unless otherwise indicated. \**P* < 0.05, \*\**P* < 0.01, \*\*\**P* < 0.001, \*\*\*\**P* < 0.0001 using Welch’s *t* test (D, F and G) and two-way ANOVA with Tukey (B) or Šídák correction (C). Error bars indicate the s.e.m.

### PGC1ɑ is essential for enhanced memory differentiation and effector functions of *PRDM1*-deficient CAR T cells

We next sought to identify upstream regulators of these mitochondrial and metabolic changes. PGC1α emerged as a potential candidate given its established roles in mitochondrial biogenesis, metabolism, and redox homeostasis, together with its increased protein abundance (**Figure 6A**) and inferred activity (**Figure 6B**) following *PRDM1* deletion. These observations led us to hypothesize that increased PGC1α activity plays a critical role in the enhanced persistence and efficacy of *PRDM1* KO CAR T cells. To test this hypothesis, PGC1α was knocked down (KD) in CAR T cells through lentiviral shRNA transduction following genetic deletion of *PRDM1* (**Figure 6C**). During *in vitro* restimulation, PGC1α KD markedly diminished the proliferative capacity (**Figure 6D**) and memory differentiation of *PRDM1* KO CAR T cells (**Figures 6E and 6F**), accompanied by an increased frequency of cells expressing the co-inhibitory receptors, PD1 and TIM3 (**Figure 6G**). To validate these findings *in vivo*, we sorted and infused CAR T cells expressing PGC1α-targeting shRNA into OCI-Ly8 xenografted mice. In line with our *in vitro* data, PGC1α KD significantly reduced expansion of CAR T cells despite *PRDM1* deletion (**Figures 6H and S5A**). More importantly, although PGC1α KD in CTRL CAR T cells still significantly prolonged the survival of tumor-bearing mice compared to the untreated group, it completely abrogated the therapeutic efficacy of *PRDM1* KO CAR T cells (**Figures 6I and S5B**). Collectively, these findings reveal that the improved functionality observed in *PRDM1*-deficient CAR T cells is mediated, at least in part, by functional engagement of PGC1α.

**Figure 6.**
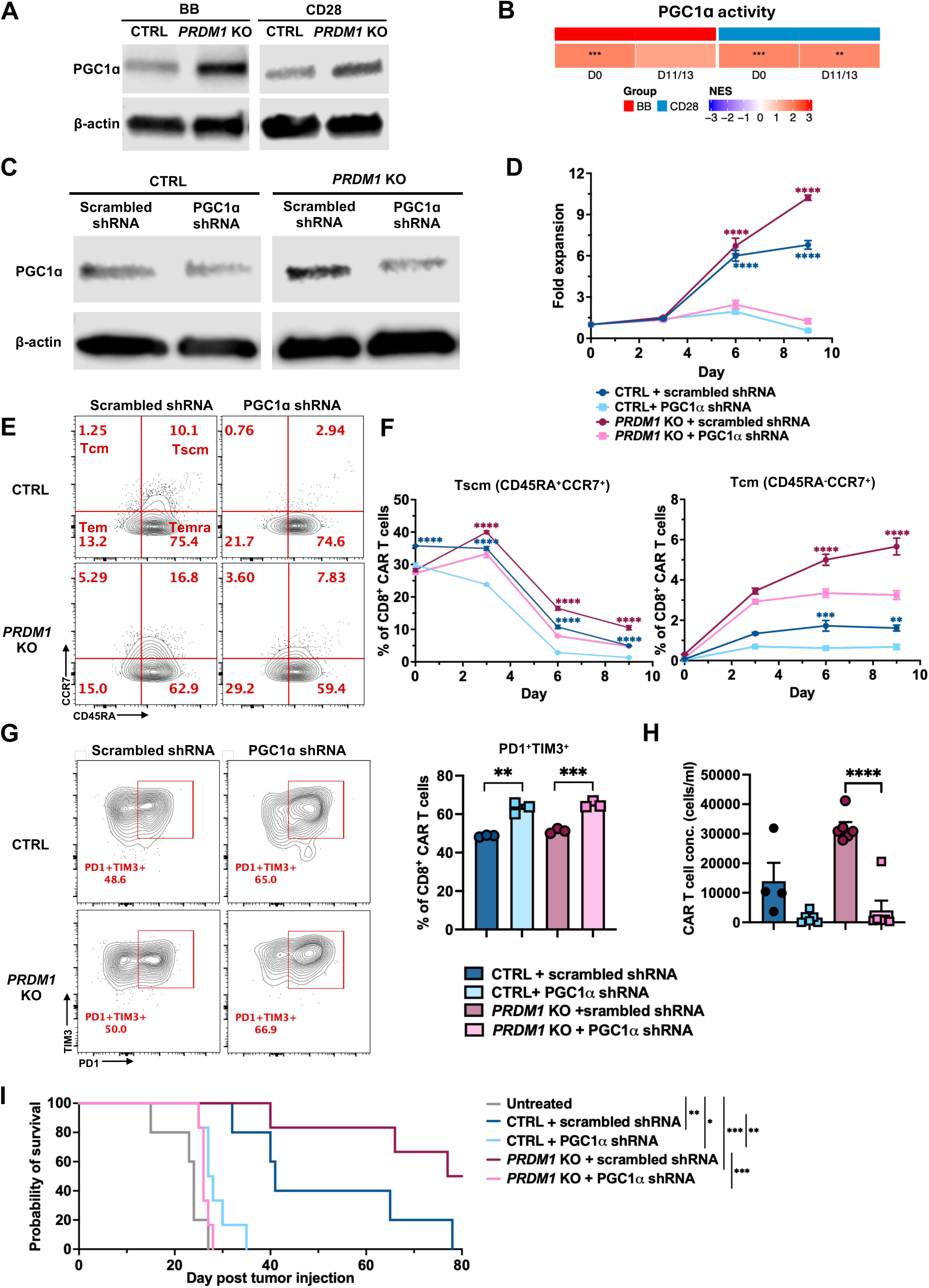
*PRDM1* deletion enhances CAR T cell memory differentiation and effector function in a PGC1α-dependent manner. **(A)** Immunoblot of PGC1α expression in CTRL and *PRDM1* KO CAR T cells, with β-actin as the loading control. **(B)** GSEA of *PRDM1* KO versus CTRL CD8^+^ CAR T cells for gene signatures associated with PGC1α activity at the indicated time points. CD8^+^ CAR T cells manufactured from 3 healthy donors were sorted before and at the end of coculture with OCI-Ly8 cells for bulk RNA-seq. NES, normalized enrichment score. *P* values were adjusted using the Benjamini–Hochberg method: \**P_adj* < 0.05, \*\**P_adj* < 0.01, \*\*\**P_adj* < 0.001. **(C)** Immunoblot of PGC1α expression in CTRL and *PRDM1* KO CAR T cells transduced with lentiviral vectors expressing scrambled control shRNA or PGC1α-targeting shRNA. β-actin served as the loading control. **(D)** Expansion dynamics of CTRL and *PRDM1* KO CAR T cells expressing scrambled control shRNA or PGC1α-targeting shRNA during restimulation with OCI-Ly8 cells, as described in Figure 2A. Statistical comparisons between PGC1α knockdown and scrambled control conditions were shown within the CTRL and *PRDM1* KO groups. **(E)** Representative flow cytometry plots showing the distribution of CD8⁺ CAR T cell subsets defined by CD45RA and CCR7 expression. Tscm (CD45RA^+^CCR7^+^), stem-cell like memory T cells; Tcm (CD45RA^-^ CCR7^+^), central memory T cells; Tem (CD45RA^-^CCR7^-^), effector memory T cells; Temra (CD45RA^+^CCR7^-^), effector memory T cells positive for CD45RA. **(F)** Frequencies of Tscm (left) and Tcm (right) subsets among CD8^+^ CAR T cells. **(G)** Representative flow cytometry plots (left) and frequencies (right) of CD8^+^ CAR T cells co-expressing PD1 and TIM3 after 8–10 days of restimulation with OCI-Ly8 cells. **(H)** Concentration of shRNA-expressing CAR T cells analyzed 12 days post infusion (n = 4–6 per group). The OCI-Ly8 xenograft experiment was performed as described in Figure 3A. **(I)** Kaplan-Meier survival curves following OCI-Ly8 transplantation (n = 4–6 per group). Data are representative of at least 2 independent experiments using CAR T cells manufactured from different healthy donors, with 3–4 technical replicates per experiment unless otherwise indicated. \**P* < 0.05, \*\**P* < 0.01, \*\*\**P* < 0.001, \*\*\*\**P* < 0.0001 using two-way ANOVA with Tukey correction for multiple comparisons (D and F), two-tailed Welch’s *t* test (G and H), and the log-rank (Mantel–Cox) test for survival analysis (I). Error bars indicate the s.e.m.

### The PPAR–PGC1ɑ agonist, bezafibrate, enhances CAR T cell performance to control hematologic and solid tumors

Our mechanistic findings identified PGC1α as an essential effector downstream of *PRDM1* loss, providing a rationale for pharmacologic intervention. We therefore investigated bezafibrate, a clinically used PPAR agonist that stimulates the PPAR–PGC1α transcriptional program. Our goal was to reproduce the therapeutic benefits of *PRDM1* deletion without further gene editing. Avoiding re-engineering could simplify manufacturing, limit editing-related genomic risks, and reduce the associated regulatory burden, facilitating broader application across CAR T cell products. We first demonstrated that bezafibrate treatment notably increased PGC1α protein levels in CAR T cells (**Figure 7A**). In lymphoma xenograft models (**Figure 7B**), bezafibrate promoted significant survival benefits in mice receiving CAR T cells containing either a BB or CD28 costimulatory domain (**Figures 7C, 7D, S6A and S6B**). Notably, bezafibrate alone exhibited no direct tumoricidal effects, indicating that bezafibrate controls tumor growth primarily through modulating CAR T cell functions. Although bezafibrate did not expedite tumor growth in our study, PGC1ɑ has shown oncogenic capacity in certain cancer types.^36–41^ We therefore evaluated whether transient exposure to bezafibrate during CAR T cell manufacturing could achieve durable therapeutic effects while mitigating potential risks associated with systemic administration (**Figure 7E**). Treatment of both BB- and CD28-based CAR T cells with bezafibrate preconditioning resulted in increased expression of canonical PGC1α target genes (**Figure 7F**) and significantly improved survival outcomes in lymphoma-grafted mice (**Figures 7G, 7H and S6C**). Further, in breast tumor-bearing mice, bezafibrate-pretreated CAR T cells showed enhanced *in vivo* expansion (**Figures 7I and 7J**), paralleled by earlier tumor regression and superior overall tumor control despite donor-to-donor variability in baseline CAR T cell potency (**Figures 7K, 7L and S6D–S6G**). These findings underscore the translational significance of bezafibrate as a safe and efficacious approach to generalize application of CAR T cell therapy.

**Figure 7.**
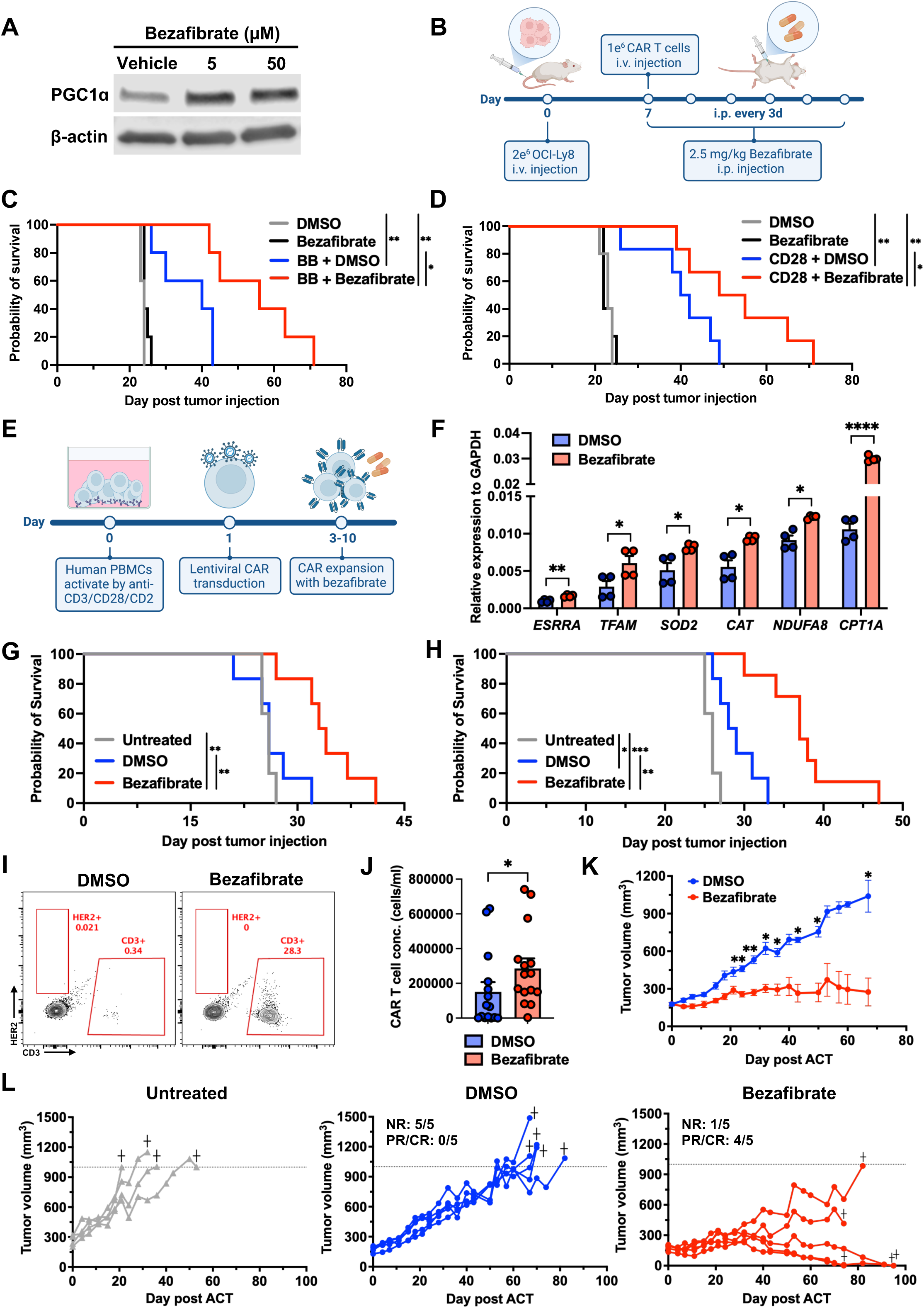
Bezafibrate synergizes the antitumor activity of CAR T cells in both hematologic and solid tumor models. **(A)** Immunoblot of PGC1ɑ expression in CAR T cells cultured *in vitro* with bezafibrate at 5 and 50 μM. DMSO-treated cells served as vehicle controls. **(B)** Experimental design of combination therapy with bezafibrate and CAR T cells. NSG mice were intravenously injected with 2 × 10^6^ OCI-Ly8 cells, followed 1 week later by infusion of 1 × 10⁶ CAR T cells via the tail vein. Bezafibrate (2.5 mg/kg) was administered intraperitoneally (i.p.) every 3 days, beginning on the day of CAR T-cell infusion and continuing until the study endpoint. DMSO served as the vehicle control. **(C)** Kaplan-Meier survival curves following OCI-Ly8 transplantation in mice receiving bezafibrate or DMSO, with or without BB-based CAR T cells (n = 4–6 mice per group). **(D)** Kaplan-Meier survival curves following OCI-Ly8 transplantation in mice receiving bezafibrate or DMSO, with or without CD28-based CAR T cells (n = 4–6 per group). **(E)** Generation of bezafibrate-preconditioned CAR T cells. Human PBMCs were activated for 1 day by T cell activators targeting CD3, CD28 and CD2, followed by transduction with CAR-encoding lentiviral vectors. Bezafibrate (30 μM) was added after transduction and replenished every 2–3 days during expansion. CAR T cells cultured with DMSO served as vehicle controls. **(F)** Expression of canonical PGC1ɑ target genes in BB- and CD28-based CAR T cells generated in the presence of bezafibrate or DMSO. Transcript levels were quantified by RT-qPCR, with each sample analyzed in technical duplicates. **(G and H)** Kaplan–Meier survival curves following OCI-Ly8 transplantation in mice receiving BB-(G) or CD28-based (H) CAR T cells preconditioned with bezafibrate or DMSO. Mice that did not receive CAR T cells served as untreated controls. Experiments were performed as described in Figure 3A (n = 5–6 mice per group). **(I)** Representative flow cytometry plots showing the frequencies of human T cells in peripheral blood 5–6 weeks post CAR T cell infusion. Mice bearing MDA-MB-361 xenografts received bezafibrate-preconditioned or DMSO-treated CAR T cells according to the protocol described in Figure 4A. **(J)** CAR T cell concentration in peripheral blood 5–6 weeks post infusion (n = 15 per group, pooled from 3 independent experiments). **(K and L)** Mean (K) and individual (L) tumor growth curves following adoptive cell transfer (ACT; n = 4– 5 per group). The dashed line indicates a tumor volume of 1,000 mm³, above which mice were evaluated for euthanasia according to the criteria described in Figure 4A. NR, no response; PR/CR, partial or complete response. Data are representative of at least 2 independent experiments using CAR T cells manufactured from different healthy donors unless otherwise indicated. \**P* < 0.05, \*\**P* < 0.01, \*\*\**P* < 0.001, \*\*\*\**P* < 0.0001 using two-tailed Welch’s *t* test (F), Mann–Whitney test (J), two-way repeated-measures ANOVA with Geisser–Greenhouse and Šídák correction (K), and the log-rank (Mantel–Cox) test for survival analysis (C, D, G and H). Error bars indicate the s.e.m.

## DISCUSSION

CAR T cell therapy can fail even when tumor antigen recognition remains unperturbed. An increasingly recognized challenge is intrinsic CAR T cell dysfunction, reflected by limited *in vivo* expansion, persistence and antitumor activity.^4–7^ In this study, we identified *PRDM1*, which encodes the transcriptional repressor BLIMP-1, as a critical regulator of CAR T cell fate. *PRDM1* was enriched in exhausted-like tumor-infiltrating CAR T cells from patients with r/r LBCL and in pre-infusion CAR T cells from clinical non-responders. This observation places *PRDM1* at both stages of the therapeutic trajectory: before infusion, where T-cell intrinsic fitness during manufacturing shapes the final product, and after infusion, where chronic antigen stimulation and the hostile tumor microenvironment drive exhaustion.

Prior studies in mouse T cells have shown that Blimp-1 promotes terminal differentiation and limits acquisition of memory properties during acute and chronic infection.^28–31^ Mechanistically, Blimp-1 regulates genes that support proliferation, cytokine response and memory potential. For instance, Blimp-1 directly represses *Il2* and the IL-2 activator *Fos*, providing negative feedback on T cell activation and expansion.^42^ Blimp-1 also targets *Il2ra* and *Cd27* by recruiting repressive histone-modifiers HDAC2 and G9a to these loci, thereby controlling cytokine responsiveness and memory-precursor formation.^43^ In addition, Blimp-1 represses the DNA-binding inhibitor *Id3*, establishing a Blimp-1–ID3–E2A axis that restricts survival and memory differentiation of effector CD8^+^ T cells.^44^ More recent work further identified a regulatory role of Blimp-1 in expression of transcription factors that promote CD8^+^ T cell stemness, including Tcf1 and Bcl6. ^45,46^

These studies provide a plausible basis for the observation of increased *PRDM1* in non-responder infusion products and exhaustion-skewed tumor-infiltrating CAR T cells with reduced proliferative capacity and memory reserve. During manufacturing, early *PRDM1* enrichment may prime CAR T cells with a lower threshold to rapidly re-engage terminal differentiation and exhaustion programs after tumor encounter, possibly accelerating the transition of CAR T cells toward short-lived effectors and dysfunctional lineages. The differential *PRDM1* induction could arise from patient heterogeneity in PBMC composition, quality, and intrinsic sensitivity to T cell activation despite standardized *ex vivo* manufacturing conditions.^47–51^ Following infusion, *PRDM1* may be further upregulated and drive exhaustion occurring during antigen exposure, hypoxia, and metabolic stress in the tumor environment.^28,52^ These observations support the hypothesis that targeting *PRDM1* during manufacturing may enhance CAR T cell efficacy by preserving memory fitness and delaying exhaustion, thereby improving *in vivo* expansion and persistence.

Recent studies have evaluated the therapeutic potential of *PRDM1* deletion in CAR T cells.^53–55^ Yoshikawa et al. showed that targeting *PRDM1* improved proliferative capacity, memory differentiation, and antitumor activity.^53^ They further demonstrated that chronically stimulated CAR T cells could partially regain memory-associated properties following *PRDM1* knockout, suggesting a role for *PRDM1* in enforcing late-stage differentiation. Jung et al. similarly reported that *PRDM1* deletion promoted stem-like features, but also identified NR4A3 induction as a compensatory mechanism that limited *in vivo* efficacy.^54^ In their prostate cancer model, dual *PRDM1* and NR4A3 ablation was required for optimal tumor control. However, this NR4A-family-related circuit has not been consistently observed across other relevant studies.^53,55^ Our work also showed that single *PRDM1* knockout was sufficient to enhance CAR T cell efficacy in both hematologic and solid tumor models despite donor heterogeneity. These differences suggest that response to *PRDM1* loss may not be universal but depend on tumor context, antigen exposure, CAR architectures, and experimental design. While multi-gene editing might improve CAR T cell performance in selected settings, it also increases manufacturing and regulatory complexities. Therefore, understanding the downstream mechanisms that mediate the durable benefits of *PRDM1* deletion is crucial for clinical translation.

Given that 4-1BB- and CD28-based CAR T cells engage distinct molecular programs after chronic stimulation,^56^ we performed bulk RNA-seq on repeatedly stimulated CAR T cells carrying either costimulatory domain to define mechanisms shared across clinically relevant CAR constructs. Notably, our analysis revealed a convergent PGC1α-associated program in *PRDM1*-deficient CAR T cells, accompanied by coordinated enrichment of mitochondrial, metabolic, and antioxidant pathways. PGC1α is a transcriptional co-activator and key mediator of mitochondrial fitness, metabolic adaptation, and effector function in CD8^+^ T cells.^57–59^ Prior studies showed that tumor-infiltrating T cells underwent mitochondrial depolarization and exhaustion in part through PGC1α repression,^57^ while enforced PGC1α expression restored mitochondrial biogenesis, metabolic reserve, and antitumor responses.^58–60^ PGC1α gain-of-function was also shown to metabolically enhance human CAR T cell activity against solid tumors.^61^ Scharping et al. further identified Blimp-1 as a negative regulator of PGC1α, showing that Blimp-

1 suppressed PGC1α promoter activity while conditional *PRDM1* deletion upregulated PGC1α expression and reduced mitochondrial oxidative stress in exhaustion-prone T cells.^52^ These findings support our model in which increased PGC1α activity contributes to enhanced mitochondrial performance and therapeutic efficacy of *PRDM1*-deficient CAR T cells. Our PGC1α knockdown experiments further implicate the functional importance of this *PRDM1*–*PGC1A* axis, as depleting PGC1α attenuated the improved expansion, memory fitness, and antitumor immunity of CAR T cells despite *PRDM1* knockout. More broadly, our mechanistic studies link *PRDM1*-driven differentiation to repression of PGC1α-dependent T cell fitness, offering a possible explanation for increased mitochondrial and metabolic resilience in early memory T cells compared with those in more differentiated or exhausted states.

To translate these insights, we examined the extent to which pharmacologic induction of the PGC1α program would reproduce the benefits of *PRDM1* deletion. Bezafibrate is a pan-PPAR agonist that activates PPARα, PPARβ/δ, and PPARγ,^62,63^ enabling these nuclear receptors to recruit coactivators such as PGC1α and regulate lipid metabolism, mitochondrial activity, and energy homeostasis.^64–67^ The long clinical history of bezafibrate as a lipid-lowering agent makes it a practical candidate for translational investigation.^68,69^ In our study, bezafibrate induced PGC1α and canonical downstream target genes in CAR T cells. Systemic bezafibrate co-treatment improved *in vivo* efficacy by enhancing CAR T cell expansion, persistence, and antitumor function, without evidence of direct tumoricidal activity. More importantly, transient exposure to bezafibrate during manufacturing was sufficient to confer durable antitumor immunity across hematologic and solid tumor models. This preconditioning approach avoids context-dependent risks on the tumor microenvironment and host tissues, offering a non-permanent and more controlled alternative to genetic modification.

In summary, our work defines molecular and functional impacts of *PRDM1* deletion across major clinical CAR platforms, establishes a *PRDM1*–*PGC1A* axis linking differentiation state to mitochondrial performance, and translates this mechanism into a manufacturing-compatible pharmacologic strategy. Future studies should investigate this modality in immunocompetent and humanized models to assess the complex effects of host immunity, tumor environmental immune suppression and treatment-related toxicities. It is also important to determine the durability of bezafibrate-induced metabolic imprinting and to evaluate the therapeutic potential of other more selective PPAR–PGC1α modulators. In parallel, mapping how *PRDM1* cooperates with broader transcriptional networks governing differentiation, exhaustion, and mitochondrial activity may refine our understanding of T cell fate development and reveal additional therapeutic opportunities to advance cellular immunotherapies.

### Limitations of the study

In this study, we identified PGC1α as an essential contributor to the enhanced fitness of *PRDM1*-deficient CAR T cells. However, the extent of preferential dependence relative to controls and the specific contribution of PGC1α upregulation remain to be clarified. Our ability to address these questions is limited by the precision of available gene engineering and protein detection methods. Advanced techniques would enable graded depletion, controlled ectopic expression, and accurate quantification of PGC1α to characterize its functional effects at finer resolution. Examining the broader relevance of these mechanistic findings would require patient-derived cells and immunocompetent or humanized models. The potential of *PRDM1* as a predictive biomarker likewise warrants further investigation. Our retrospective analyses linked its expression to infusion product quality and clinical outcomes. Independent cohorts, adjustment for relevant covariates, and prospective studies are needed to determine whether *PRDM1* expression could add prognostic value beyond established factors and inform treatment decisions.

## METHODS AND MATERIALS

### Cell lines

MDA-MB-361 and tdTomato-expressing OCI-Ly8 cells were cultured at 37°C in a humidified incubator with 5% CO₂ in RPMI-1640 (Gibco) supplemented with 10% (v/v) FBS, 1% (v/v) penicillin– streptomycin, 1% (v/v) MEM non-essential amino acids (Gibco), 1% (v/v) HEPES (Gibco), 1% (v/v) GlutaMAX (Gibco), and 1% (v/v) sodium pyruvate (Gibco). Adherent MDA-MB-361 cells were maintained at 40 to 80% confluence; suspension OCI-Ly8 cells were passaged every 2–3 days to maintain a density of 0.2 to 1.0 × 10^6^ cells/ml. tdTomato-expressing OCI-Ly8 cells were kindly provided by the Huang laboratory (University of Chicago, USA), and MDA-MB-361 cells were kindly provided by the Greene laboratory (University of Chicago, USA). All cell lines were routinely tested and confirmed negative for mycoplasma contamination using the PCR-based assay (Applied Biological Materials).

### Human primary cells

Healthy donor peripheral blood mononuclear cells (PBMCs) were purchased from Lonza or isolated by Ficoll density-gradient centrifugation (Sigma-Aldrich) from whole blood collected through the University of Chicago Medicine Blood Donation Center. The protocols were approved by the University of Chicago Institutional Review Board. Freshly isolated PBMCs were cryopreserved in fetal bovine serum (FBS; GeminiBio) containing 10% (v/v) dimethyl sulfoxide (DMSO; Thermo Fisher Scientific) and stored in liquid nitrogen until use. Commercial PBMCs (Lonza) were handled according to the manufacturer’s instructions. Before experiments, cryopreserved PBMCs were thawed and cultured in complete T cell medium (TexMACS™ GMP Medium [Miltenyi Biotec] supplemented with 10% [v/v] FBS, 1% [v/v] penicillin-streptomycin [Gibco], 5 ng/ml recombinant human IL-7 and IL-15 [PeproTech]) at 37°C in a humidified incubator with 5% CO₂.

### Design and vector cloning of lentiviral constructs

CAR constructs were cloned into self-inactivating (SIN) pHR lentiviral transfer vectors for constitutive expression. Anti-CD19 CARs comprised an FMC63-derived single-chain variable fragment (scFv), either a 4-1BB or CD28 costimulatory domain, and a CD3ζ signaling domain, with a directly fused monomeric enhanced green fluorescent protein (mEGFP) or blue fluorescent protein (BFP) reporter. The anti-HER2 CAR comprised a 4D5-derived scFv, a 4-1BB costimulatory domain, and a CD3ζ signaling domain and was fused to mEGFP. An shRNA targeting *PPARGC1A*, which encodes PGC1α, and a non-targeting control shRNA were cloned into BFP-expressing, SIN pLKO.1 lentiviral transfer vectors. The full shRNA sequences were as follows: *PPARGC1A* shRNA, forward, 5′-CCGGGACTATTGCCAGTCAATTAATCTCGAGATTAATTGACTGGCAATAGTCTTTTTG-3′, and reverse, 5′-AATTCAAAAAGACTATTGCCAGTCAATTAATCTCGAGATTAATTGACTGGCAATAGTC-3′; non-targeting control shRNA, forward, 5′-CCGGCAACAAGATGAAGAGCACCAACTCGAGTTGGTGCTCTTCATCTTGTTGTTTTTG-3′, and reverse, 5′-AATTCAAAAACAACAAGATGAAGAGCACCAACTCGAGTTGGTGCTCTTCATCTTGTTG-3′. All constructs were generated using standard restriction enzyme digestion and ligation protocols with reagents from New England Biolabs (NEB). Plasmids were propagated in NEB Stable Competent Escherichia coli and purified using the ZymoPURE II Plasmid Midiprep Kit (Zymo Research). Construct sequences were verified by Sanger sequencing at the Sanger (DNA) Sequencing Core (University of Chicago).

### Lentivirus production

Lentiviral vectors were produced using Lenti-X 293T cells (Takara Bio) maintained in Dulbecco’s Modified Eagle Medium (DMEM; Gibco) supplemented with 10% FBS at 37°C. Cells were co-transfected with transfer plasmids, pMD2.G VSV-G envelope plasmids (Addgene), and pCMV-dR8.2 dvpr packaging plasmids (Addgene) using polyethylenimine (PEI; Polysciences). Plasmids and PEI were premixed in Opti-MEM I Reduced Serum Medium (Gibco) before addition to Lenti-X 293T cells. Two days after transfection, virus-containing culture supernatants were collected, clarified by centrifugation, passed through a 0.45-μm filter, and concentrated by PEG precipitation using a solution containing 40% (w/v) PEG-8000 (Fisher Scientific) and 1.2 M NaCl (Sigma-Aldrich) in PBS.

### Generation of CAR T cells

Human PBMCs were activated by ImmunoCult Human CD3/CD28/CD2 T Cell Activator (STEMCELL Technologies) per manufacturer’s instructions. One day following stimulation, cells were transduced with CAR-encoding lentivirus in the presence of 10 μg/ml protamine sulfate (Sigma-Aldrich) and expanded in complete T cell medium for 10 to 13 days before downstream experiments. To generate bezafibrate-preconditioned CAR T cells, bezafibrate (Santa Cruz Biotechnology) was prepared as a 50 mg/ml stock solution in DMSO, added at 30 μM immediately after transduction and replenished every 3 days throughout the culture period. Vehicle-control cells received an equivalent volume of DMSO. To deplete PGC1α, cells were co-transduced with lentiviral vectors encoding the CAR and a *PPARGC1A*-targeting or non-targeting control shRNA. PGC1α knockdown was confirmed by immunoblotting.

### CRISPR/Cas9-mediated gene editing

After 24 hours of T cell activation and before lentiviral transduction, electroporation was performed using the Neon Transfection System (Thermo Fisher Scientific) according to the manufacturer’s instructions. In brief, three synthetic single-guide RNAs (sgRNAs; Synthego) targeting multiple sites of *PRDM1* were pooled and mixed with recombinant SpCas9 protein (QB3 MacroLab, University of California, Berkeley) at a 5:2 total sgRNA-to-Cas9 molar ratio. For each electroporation, 50 pmol of each sgRNA (150 pmol total) was combined with 60 pmol SpCas9 and incubated for 10 min at room temperature to form ribonucleoprotein (RNP) complexes. A total of 5 × 10^5^ cells were resuspended in 10 μL Neon Resuspension Buffer T and electroporated with the RNP complexes using a 10-μL Neon tip at 1,400 V, a pulse width of 10 milliseconds, and 3 pulses. Control cells received RNP complexes assembled with an equivalent amount of Negative Control SpCas9 sgRNA #1 (Synthego) and SpCas9 protein. Following electroporation, cells were transduced with CAR-encoding lentivirus and expanded as described above. Loss of BLIMP-1 encoded by *PRDM1* was confirmed by immunoblotting and functional assays. The *PRDM1*-targeting sgRNA sequences were as follows: target 1, 5′-CATCAGCACCAGAATCCCAG-3′; target 2, 5′-CGGATGGGGTAAACGACCCG-3′; and target 3, 5′-TCGATGACTTTAGAAGACGT-3′.

### In vitro CAR T cell restimulation assay

CAR T cells were serially challenged with human OCI-Ly8 DLBCL cells every 2–3 days at an effector-to-target (E:T) ratio ranging from 1:8 to 1:16 optimized for each donor in R10 medium (RPMI-1640 supplemented with 10% [v/v] FBS, 1% [v/v] penicillin–streptomycin, 1% [v/v] HEPES (Gibco), and 1% [v/v] GlutaMAX). At each time point, cells were enumerated using fluorescent counting beads (Spherotech) and analyzed by flow cytometry to assess CAR T cell expansion, differentiation and effector functions. The remaining cells were resuspended in a 1:1 mixture of conditioned (old) and fresh R10 medium, with additional OCI-Ly8 cells for continued challenge. This process was repeated for 10–15 days for total 4–5 rounds of stimulation until CAR T cell fold-change fell below 1 or tumor cell fold-change exceeded 1, which served as operational criteria for CAR T-cell dysfunction.

### Seahorse assay

CD8^+^ CAR T cells were sorted using FACSymphony S6 Cell Sorter (BD Biosciences) and plated on Seahorse XFe96 culture microplates coated with Cell-Tak (Corning) at 150,000 cells per well in Seahorse XF DMEM medium supplemented with 10 mM glucose, 1 mM pyruvate, and 2 mM glutamine (Agilent Technologies). In mito stress tests, oligomycin (2 μM), FCCP (1 μM) and rotenone/antimycin A (0.5 μM) from the Seahorse XF Cell Mito Stress Test Kit (Agilent Technologies) were injected sequentially to measure the changes in cellular metabolic activity.

### Flow cytometry

For routine cell-surface staining, cells were first incubated with Fc-blocking reagent for 5 min at room temperature, followed by staining with fluorophore-conjugated anti-human antibodies (Table S1) in FACS buffer (PBS supplemented with 2% [v/v] FBS) for 30 min at 4°C. Chemokine receptor CCR7 staining was performed separately for 15 min at 37°C before staining for other markers. Dead cells were excluded using the LIVE/DEAD Fixable Near-IR Dead Cell Stain Kit (Invitrogen). For mitochondrial analyses, cells were incubated with MitoSOX (Invitrogen) and MitoTracker dyes (MitoTracker Green and MitoTracker Deep Red; Invitrogen) diluted in Hanks’ balanced salt solution (HBSS; Gibco) for 30 min at 37°C. For intracellular staining, cells were fixed and permeabilized using the eBioscience Foxp3/Transcription Factor Staining Buffer Set (Invitrogen) according to the manufacturer’s instructions. Flow cytometric data were acquired using the Aurora (Cytek Biosciences) or NovoCyte Penteon 5-30 (Agilent Technologies) flow cytometer and analyzed using FlowJo v10.10.0 (BD Biosciences).

### Quantitative reverse-transcription PCR (RT-qPCR)

Total RNA was isolated from the indicated cells using the Direct-zol RNA Miniprep Kit (Zymo Research) and reverse-transcribed using High-Capacity cDNA Reverse Transcription Kits (Applied Biosystems) per manufacturer’s instructions. RT-qPCR was performed using PowerUp SYBR Green Master Mix on a ViiA 7 Real-Time PCR System (Applied Biosystems). Relative gene expression was normalized to housekeeping gene GAPDH. Primer sequences were provided in Table S2.

### Bulk RNA sequencing

For RNA isolation, CD8⁺ anti-CD19 CAR T cells were collected before stimulation and after the fourth or fifth tumor challenge. Pre-stimulation samples were sorted directly, whereas post-challenge samples were first enriched by depleting CD19-expressing OCI-Ly8 cells using the MojoSort Human CD19 Selection Kit (BioLegend). Sorting was performed using a FACSymphony S6 Cell Sorter with antibodies listed in Table S1. Total RNA was extracted from sorted cells using the Direct-zol RNA Miniprep Kit. RNAseq libraries were prepared using the KAPA Hyper Stranded mRNA Library Kit (Roche), pooled, and quantified by qPCR before sequencing at the Roy J. Carver Biotechnology Center (University of Illinois Urbana-Champaign). Paired-end sequencing (150 bp per end) was then performed on the NovaSeq X Plus (Illumina) using v1.0 sequencing kits and a single lane of a 10B flow cell. Bioinformatic analysis was conducted by the Center for Research Informatics (University of Chicago). Raw reads were processed using nf-core/rnaseq, with FastQC for read quality assessment, Trim Galore for adapter and quality trimming, and MultiQC for reporting quality metrics. Reads were aligned to the human reference genome (GRCh38) using STAR, and transcript abundances were quantified using Salmon and summarized into gene-level counts using tximport. Lowly expressed genes were filtered based on counts per million (CPM), and protein-coding genes were retained for downstream analysis. Counts were normalized using the trimmed mean of M-values (TMM) method in edgeR. For gene set enrichment analysis (GSEA), genes were ranked by sign(log₂ fold change) × −log₁₀(P value), excluding genes with missing or nonfinite ranking values. Gene sets were obtained from MSigDB using msigdbr and included Hallmark, KEGG, Reactome, and Gene Ontology Biological Process (GOBP) with pathways related to mitochondrial function, metabolism, and redox homeostasis. A custom PGC1α gene signature was also evaluated by GSEA using the gene list (Table S3) curated from the published PGC1α overexpression study.^61^ The GSEA was performed using fgsea with standard scoring and gene set sizes of 5–500 genes. P values were adjusted using the Benjamini–Hochberg method. Normalized enrichment scores (NES) were visualized using ComplexHeatmap.

### Single-cell RNA sequencing

To examine *PRDM1* expression and regulatory activity in tumor-infiltrating CD8^+^ T cells, the single-cell dataset from Li et al. (EGAS50000001022),^32^ comprising 217 large B cell lymphoma (LBCL) tumor biopsies, was reanalyzed with the original cell-subset annotations. Scanpy was used for uniform manifold approximation and projection (UMAP) visualization and exhaustion scoring with sc.tl.score_genes.^70^ Differentially expressed genes (DEGs) between exhausted CD8^+^ T cells and all remaining CD8^+^ subsets were identified by the Wilcoxon rank-sum test using sc.tl.rank_genes_groups. For enhancer regulon (eRegulon) analysis, CD8^+^ T cells with paired, high-quality snRNA-seq and snATAC-seq profiles were analyzed using SCENIC+,^71^ as previously described, to identify eRegulons that capture trios of candidate transcription factors, target enhancer genomic regions, and co-regulated target genes. Subset specificity was quantified using regulon_specificity_scores. To investigate *PRDM1* expression in tumor-infiltrating CAR T cells, the scRNA-seq dataset from Arita et al. (EGAS50000001982) was reanalyzed.^33^ CD8^+^ T cells were selected from relapsed/refractory LBCL tumors in CAR T-cell–naïve patients (n = 5) and patients refractory to CAR T-cell therapy (n = 7), retaining pre-established subset annotations and CAR detection status. UMAP embeddings and transcriptomic signature scores were generated using Seurat and AddModuleScore, respectively.^72^ Differential gene expression between CAR^+^CD8^+^ T cells and all remaining CD8^+^ T cells was assessed by the Wilcoxon rank-sum test using FindMarkers. To correlate *PRDM1* expression and activity in infusion products with clinical response, scRNA-seq data from Haradhvala et al. (GSE197268) were processed and analyzed using Seurat with original cell annotations.^10^ Microarray data from Rutishauser et al. (GSE17211) were analyzed to evaluate differential gene expression using linear models in limma.^29^ DEGs in *PRDM1* conditional-knockout cells were retained using thresholds of log2 fold change >0.5 and adjusted *P* < 0.05 (Table S4). The gene set was scored per group using AddModuleScore in Seurat. To identify pathways associated with *PRDM1* expression, patients with LBCL (GSE197268) were stratified into *PRDM1*-high (n = 3) and *PRDM1*-low (n = 5) groups based on *PRDM1* expression in their infusion products, using a 0.35 quantile-based cutoff for subsequent pre-ranked GSEA.

### Western blot analysis

Whole-cell protein lysates were prepared using Cell Lysis Buffer (Cell Signaling Technology) with 1% (v/v) Halt Protease and Phosphatase Inhibitor Cocktail (Thermo Fisher Scientific). An equal amount of protein (15-20 μg) was resolved by SDS–PAGE using 4-20% Mini-PROTEAN TGX Precast Gels (Bio-Rad) and transferred onto Immobilon-FL PVDF membranes (Merck Millipore). The membranes were blocked with 5% bovine serum albumin (BSA; Fisher Scientific) in Tris-buffered saline containing 0.1% Tween 20 (TBST; Cell Signaling Technology) and incubated overnight at 4°C with primary antibodies: rabbit anti-β-actin (1:4000; Cell Signaling Technology), mouse anti-BLIMP-1 (1:1000; Novus Biologicals), or rabbit anti-PGC1α (1:1000; Cell Signaling Technology). After washing with TBST, membranes were incubated for 1 hour at room temperature with HRP-conjugated goat anti-mouse or goat anti-rabbit IgG secondary antibodies (1:2000; Cell Signaling Technology). Protein bands were detected using Clarity Western ECL Substrate (Bio-Rad) and visualized with a LI-COR imaging system.

### Electron microscopy

Sample preparation and imaging were performed by the Advanced Electron Microscopy Facility (University of Chicago). Briefly, repeatedly stimulated anti-CD19 CAR T cells were washed with serum-free medium and fixed at 37°C with 2% glutaraldehyde and 4% paraformaldehyde (Electron Microscopy Sciences) in 0.1 M sodium cacodylate buffer. Fixed cells were washed with 0.1 M sodium cacodylate buffer and post-fixed with 1% osmium tetroxide in 0.1 M sodium cacodylate buffer for 1 hour. Samples were subsequently stained with 1% uranyl acetate in maleate buffer for 1 hour and sequentially dehydrated in 25%, 50%, 70%, 95%, and 100% ethanol, followed by 100% propylene oxide. Samples were infiltrated with propylene oxide and Spurr resin at a 2:1 ratio for two 30-minute incubations, followed by overnight infiltration in a 1:1 mixture. Samples were then subjected to six 1-hour exchanges in 100% Spurr resin and polymerized at 60°C for 1–2 days. Ultrathin sections (90 nm) were cut using an EM UC6 ultramicrotome (Leica Microsystems) and stained with uranyl acetate and lead citrate. Images were acquired using a Tecnai F30 transmission electron microscope (FEI Company) at 300 kV and analyzed using ImageJ.

### Mouse xenograft studies

Mouse studies were performed with 8- to 12-week-old NOD.Cg-*Prkdc^scid^ Il2rg^tm1Wjl^*/SzJ mice (NSG; strain no. 005557; The Jackson Laboratory). All animal procedures were approved by the University of Chicago Institutional Animal Care and Use Committee and performed in accordance with the National Institutes of Health guidelines. For the lymphoma xenograft model, male and female NSG mice were injected intravenously with 2 × 10^6^ OCI-Ly8 cells and 7 days later, mice received 1 × 10^6^ CAR T cells via the tail vein. For bezafibrate co-treatment experiments, bezafibrate reconstituted in DMSO was diluted in PBS at 0.25 mg/ml immediately before administration and injected intraperitoneally at 2.5 mg/kg every 3 days, starting on the day of CAR infusion and continuing until the experimental endpoint. Vehicle-treated-mice received the corresponding concentration of DMSO in PBS following the same dosing schedule. Mice were monitored regularly and euthanized upon reaching protocol-defined humane endpoints associated with lymphoma progression. In the indicated lymphoma experiments, spleens were collected at the humane endpoint, and splenic CAR T cell differentiation was analyzed by flow cytometry. For the breast cancer model, 1 × 10^7^ MDA-MB-361 cells were suspended in a 1:1 mixture of ice-cold phenol red-free Matrigel (Corning) and PBS and injected orthotopically into the mammary fat pad of female NSG mice. Tumor volumes were calculated from caliper measurements every 3 to 4 days: tumor volume = (length x width x height) x π/6. When the average tumor size reached 200– 350 mm^3^, mice received 2 to 3 × 10^6^ CAR T cells intravenously. Mice were euthanized when tumors exceeded 1,500 mm^3^ or 1,000 mm^3^ with ulceration or bleeding. To monitor CAR T cell expansion and persistence, peripheral blood was collected at the indicated time points by submandibular bleeding. Red blood cells were lysed for 10 minutes at room temperature using buffer containing 140 mM NH₄Cl and 17 mM Tris-HCl before surface staining. The absolute numbers of CAR T cells were determined using fluorescent counting beads. For analysis of tumor-infiltrating CAR T cells, breast tumors were harvested 5–6 weeks post CAR T cell infusion and digested with collagenase IV (Sigma-Aldrich) and DNase I (Roche) for 30 min at 37°C. Tumor-derived cells were stimulated with 50 ng/ml phorbol 12-myristate 13-acetate (PMA; Sigma-Aldrich) and 1 μM ionomycin (Sigma-Aldrich) for 1 hour. Then, brefeldin A (GolgiPlug; BD Biosciences) and monensin (GolgiStop; BD Biosciences) were added for an additional 4 hours before intracellular cytokine staining.

### Statistical analysis

Statistical analyses were performed using Prism (GraphPad Software). Comparisons between two groups were performed using Welch’s t test or Mann–Whitney test. Comparisons involving multiple groups were performed using one-way or two-way analysis of variance (ANOVA) with Šídák or Tukey correction, according to the experimental design and data distribution. Tumor growth curves were analyzed using two-way repeated-measures ANOVA, with Geisser– Greenhouse and Šídák correction. Survival data were presented as Kaplan–Meier curves and analyzed using the log-rank (Mantel–Cox) test. Data are presented as the mean ± standard error of the mean (s.e.m.). Statistical significance is denoted as follows: \**P* < 0.05, \*\**P* < 0.01, \*\*\**P* < 0.001, and \*\*\*\**P* < 0.0001. The statistical tests used per figure are specified in the corresponding figure legends.

## ACKNOWLEDGMENTS

We would like to thank Drs. James L. Labelle, Jun Huang, Geoffrey Greene, and Marsha Rosner for sharing mice, cell lines and plasmid constructs central to this study. Special thanks to Aniruddhsingh Solanki of the University of Chicago Animal Resources Center for assistance with mouse injections. We also thank the University of Chicago Human Disease and Immune Discovery, Cytometry and Antibody Technology facility, Advanced Electron Microscopy Core Facility, and BioPhysics Core Facility for technical support and data acquisition during the project. Thanks to the University of Chicago Center for Research Informatics for bulk RNA-seq analysis. Single-cell RNA-seq datasets were generated by the MD Anderson Lymphoid Malignancies Program. We are grateful to Drs. Yuxuan Miao, Maria-Luisa Alegre, and Alexander Muir for constructive feedback and helpful discussions as thesis committee members, and to Dr. Peter Savage for insightful conversations during joint meetings. This work was funded by the Jonas Center for Cellular Therapy to J.K. S.W. was supported by the NCI T32 Multi-disciplinary Training Grant in Cancer Research (5T32CA009594-34 and 5T32CA009594-35), Robert C and Mary Jane Gallo Scholarship, and Fitch Scholarship at the University of Chicago.

## Author contributions

Conceptualization, S.W., L.X., X.C. and J.K.; methodology, S.W., X.C., M.R.G., X.L., A.E., Y.H., A.C., J.C., and L.X.; validation, S.W.; investigation, S.W.; writing, S.W. and J.K.; supervision, J.K.; funding acquisition, J.K.

