## Supplementary Figures for "Harnessing the *PRDM1–PGC1Α* axis to enhance CAR T cell efficacy"

**Figure S1. Repeated stimulation with OCI-Ly8 tumor cells induces CD19-targeted CAR T cell dysfunction.**

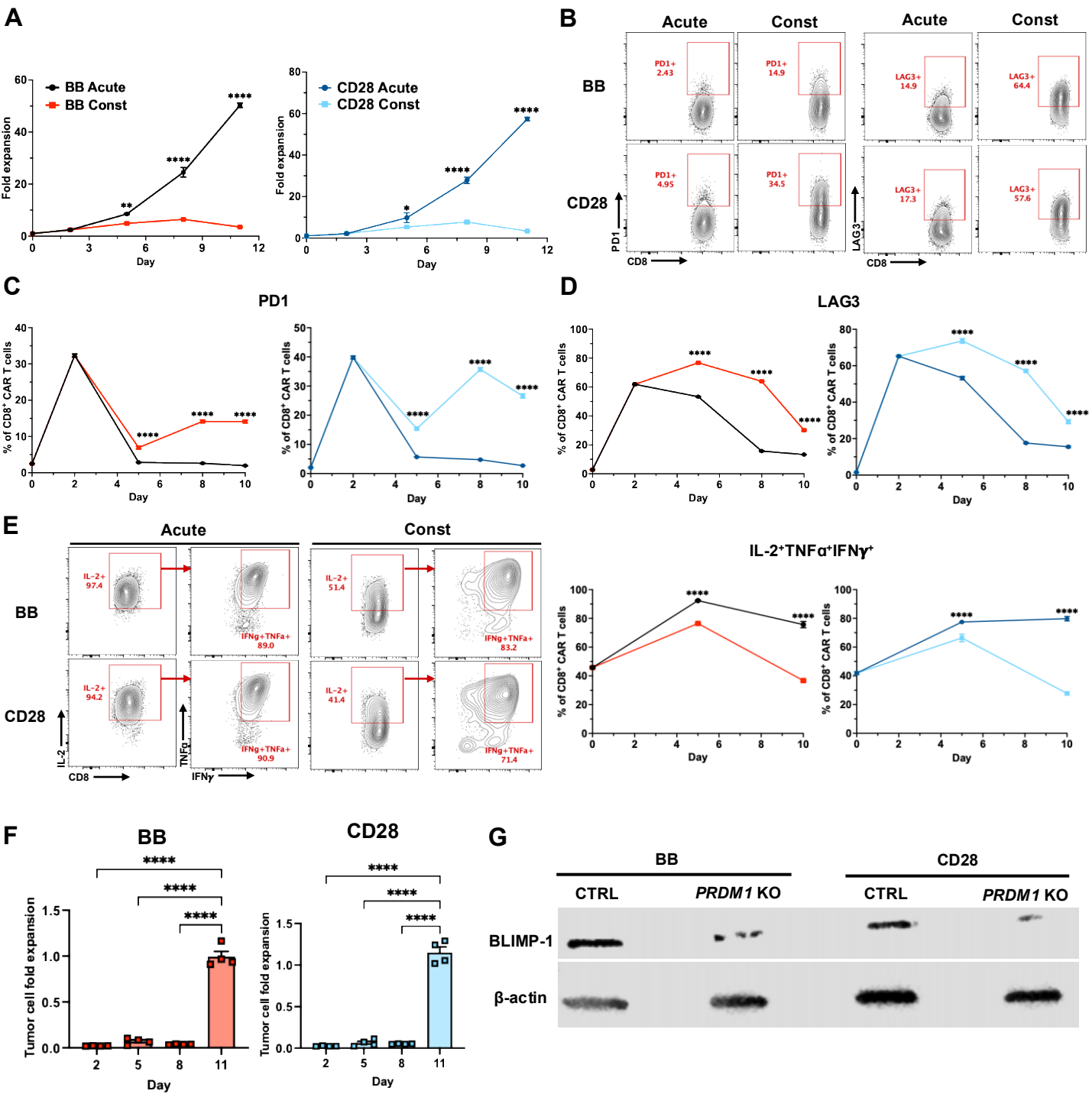

Figure S2. *PRDM1*-deficient CAR T cells prolong survival of lymphoma-xenografted NSG mice.

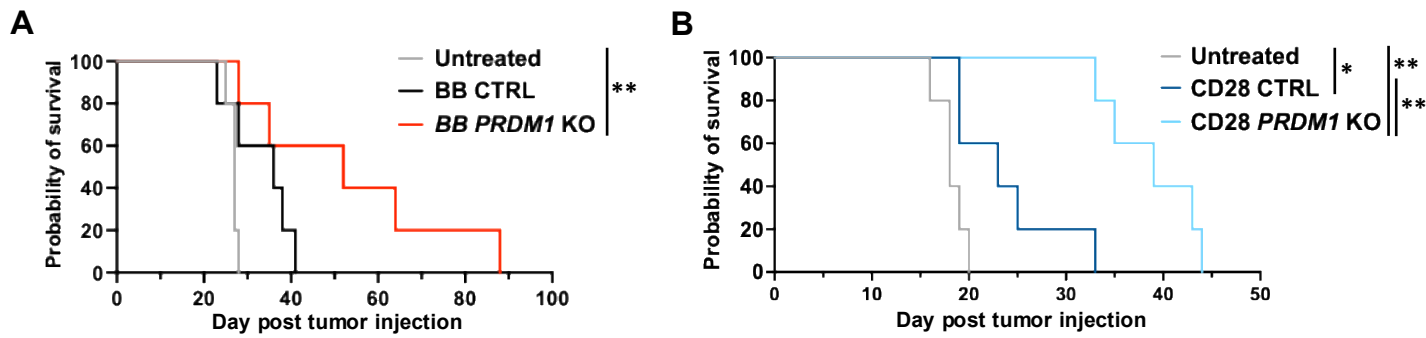

**Figure S3. *PRDM1* deletion enhances antitumor efficacy of HER2-directed CAR T cells against breast tumors.**

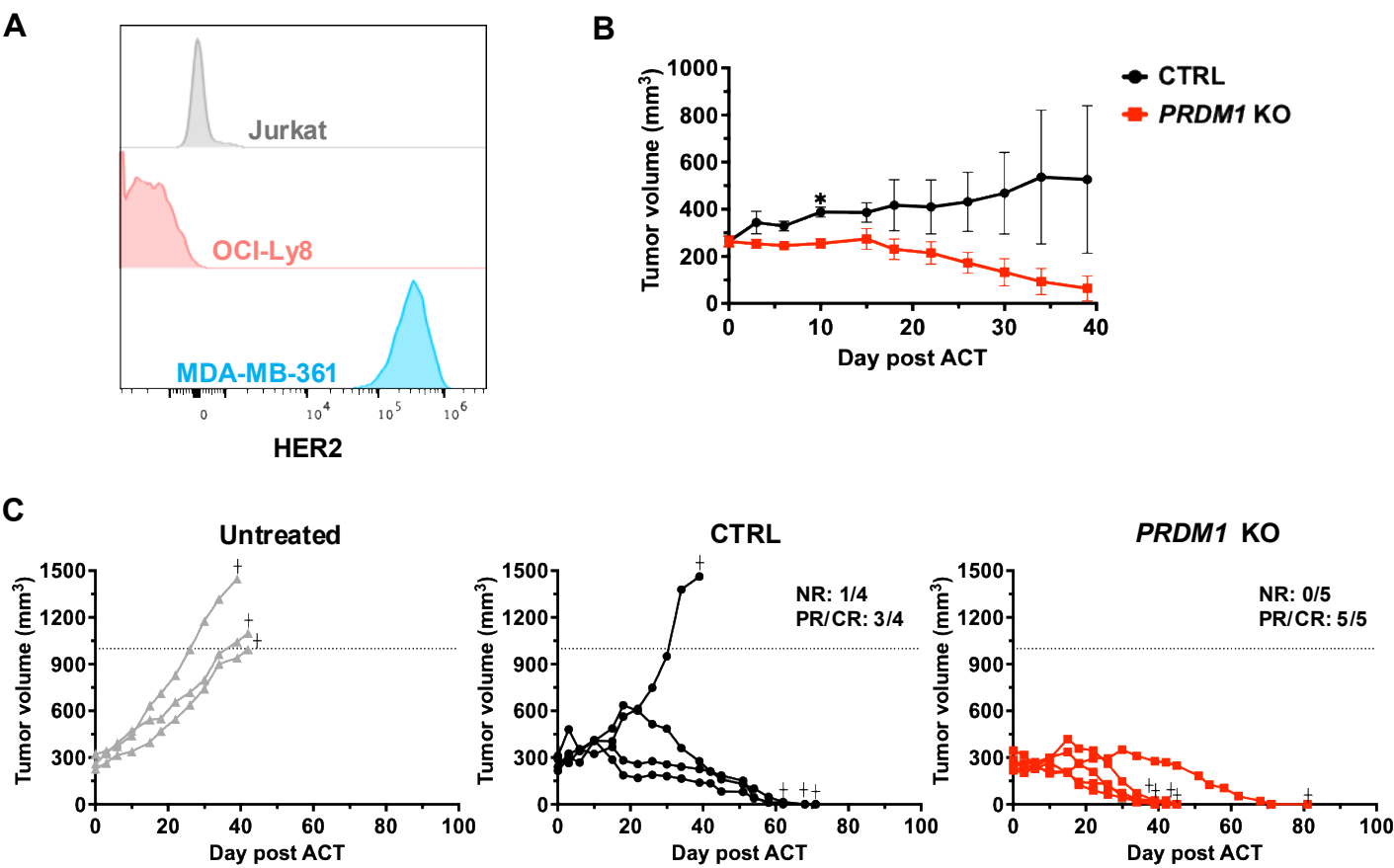

**Figure S4. *PRDM1* deletion promotes mitochondrial, metabolic, and antioxidant programs in CAR T cells.**

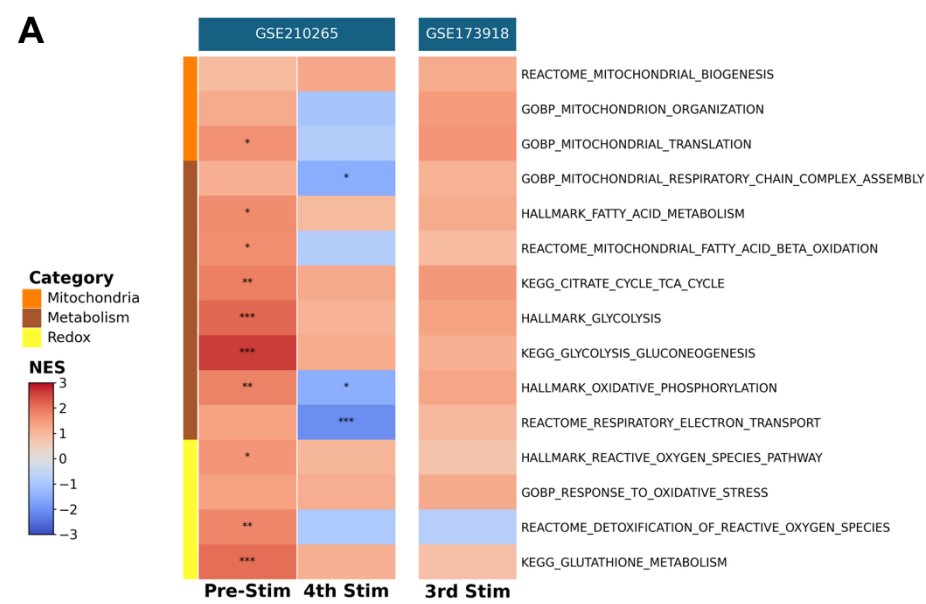

Figure S5. PGC1α is essential for enhanced antitumor activity of *PRDM1*-deficient CAR T cells

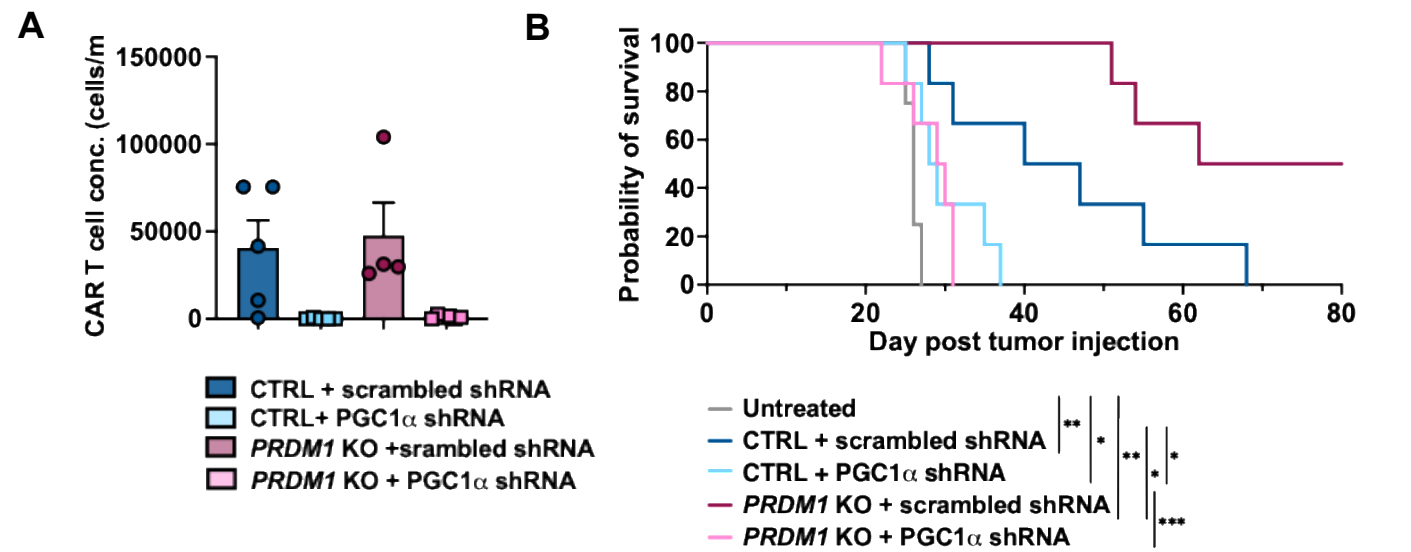

**Figure S6. Bezafibrate potentiates CAR T cell-mediated tumor control in hematologic and solid tumor models.**

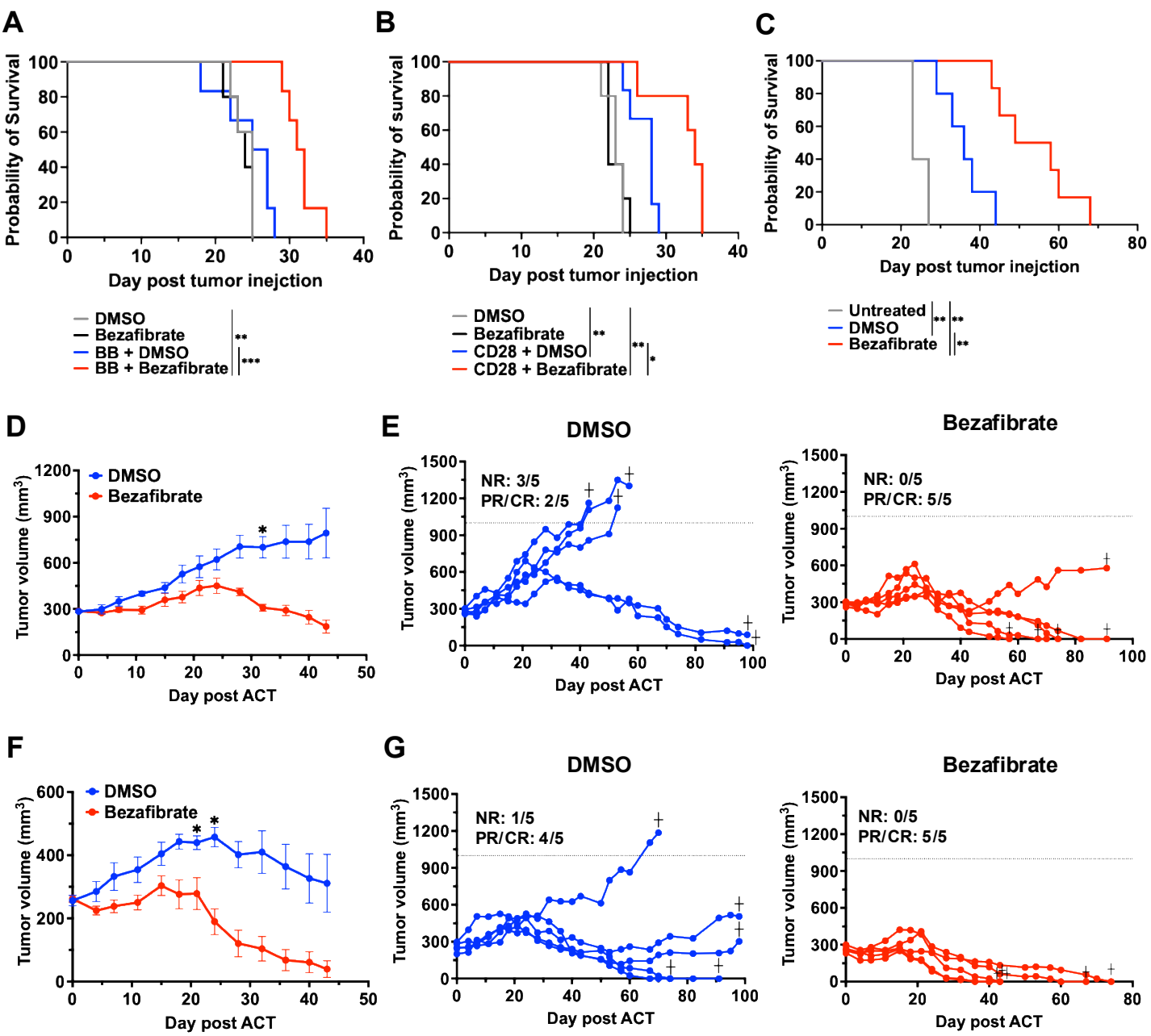
