## Supplementary Figure legends for "Harnessing the *PRDM1–PGC1Α* axis to enhance CAR T cell efficacy"

**Figure S1. Repeated stimulation with OCI-Ly8 tumor cells induces CD19-targeted CAR T cell dysfunction.**

**(A)** Expansion dynamics of BB- or CD28-based CAR T cells under acute or constant stimulation. Acutely stimulated CAR T cells (BB Acute and CD28 Acute) were stimulated once with OCI-Ly8 cells on day 0 and maintained in fresh R10 medium supplemented with 5 ng/mL recombinant human IL-7 and IL-15. Constantly stimulated CAR T cells were serially challenged with OCI-Ly8 cells as described in Fig. 2A.

**(B–D)** Representative flow cytometry plots (B) and percentages of CD8<sup>+</sup> CAR T cells expressing PD1 (C) or LAG3 (D) during restimulation, respectively.

**(E)** Representative flow cytometry plots (left) and frequencies (right) of CD8<sup>+</sup> CAR T cells producing effector cytokines, IL-2, IFNγ and TNFα. Before and during coculture, CAR T cells were restimulated with PMA/ionomycin for 5 hours in the presence of brefeldin A/monensin before analysis.

**(F)** Fold expansion of OCI-Ly8 cells during coculture with BB- or CD28-based CAR T cells under constant stimulation, calculated relative to the tumor-cell count in the preceding culture.

**(G)** Immunoblot of BLIMP-1 expression in CTRL and *PRDM1* KO CAR T cells, with β-actin as the loading control. Data are representative of at least 3 independent experiments using CAR T cells manufactured from different healthy donors, with 3–4 technical replicates per experiment. \*$P < 0.05$, \*\*$P < 0.01$, \*\*\*$P < 0.001$, \*\*\*\*$P < 0.0001$ using ordinary one-way (F) or two-way (A, C, D and E) ANOVA with Šídák correction for multiple comparisons. Error bars indicate the s.e.m.

**Figure S2. *PRDM1*-deficient CAR T cells prolong survival of lymphoma-xenografted NSG mice.**

**(A and B)** Kaplan-Meier survival curves following OCI-Ly8 transplantation (n = 5 per group). \*$P < 0.05$, \*\*$P < 0.01$, \*\*\*$P < 0.001$, \*\*\*\*$P < 0.0001$ using the log-rank (Mantel–Cox) test for survival analysis.

**Figure S3. *PRDM1* deletion enhances antitumor efficacy of HER2-directed CAR T cells against breast tumors.**

**(A)** HER2 expression in the human breast cancer cell line MDA-MB-361. The CD4<sup>+</sup> T cell line Jurkat and OCI-Ly8 cells served as negative controls.

**(B and C)** Mean (B) and individual (C) tumor growth curves following adoptive cell transfer (ACT). The dashed line indicates a tumor volume of 1,000 mm<sup>3</sup>, above which mice were evaluated for euthanasia according to the criteria described in Figure 4A. NR, no response; PR/CR, partial or complete response. Data are from an independent experiment using CAR T cells manufactured from a different healthy donor (n = 3–5 mice per group). \*$P < 0.05$, \*\*$P < 0.01$, \*\*\*$P < 0.001$, \*\*\*\*$P < 0.0001$ using two-way repeated-measures ANOVA with Geisser–Greenhouse and Šídák correction. Error bars indicate the s.e.m.

**Figure S4. *PRDM1* deletion promotes mitochondrial, metabolic, and antioxidant programs in CAR T cells.**

(A) GSEA comparing *PRDM1* KO versus CTRL CD8<sup>+</sup> CAR T cells before or at the end of tumor-cell restimulation, as indicated (GSE210265 and GSE173918). NES, normalized enrichment score. *P* values were adjusted using the Benjamini–Hochberg method: \**P*<sub>adj</sub> < 0.05, \*\**P*<sub>adj</sub> < 0.01, \*\*\**P*<sub>adj</sub> < 0.001.

**Figure S5. PGC1α is essential for enhanced antitumor activity of *PRDM1*-deficient CAR T cells.**

(A) Concentration of shRNA-expressing CAR T cells 13 days post infusion (n = 4–6 per group). The mouse experiment was conducted as described in Figure 3A.

(B) Kaplan-Meier survival curves following OCI-Ly8 transplantation (n = 4–6 per group). Data are from an independent experiment using CAR T cells manufactured from a different healthy donor. \**P* < 0.05, \*\**P* < 0.01, \*\*\**P* < 0.001, \*\*\*\**P* < 0.0001 using two-tailed Welch's *t* test (A), and the log-rank (Mantel–Cox) test for survival analysis (B). Error bars indicate the s.e.m.

**Figure S6. Bezafibrate potentiates CAR T cell-mediated tumor control in hematologic and solid tumor models.**

(A) Kaplan-Meier survival curves following OCI-Ly8 transplantation in mice receiving bezafibrate or DMSO, with or without BB-based CAR T cells (n = 4–6 mice per group).

(B) Kaplan-Meier survival curves following OCI-Ly8 transplantation in mice receiving bezafibrate or DMSO, with or without CD28-based CAR T cells (n = 4–6 per group).

(C) Kaplan–Meier survival curves following OCI-Ly8 transplantation in mice receiving CD28-based CAR T cells preconditioned with bezafibrate or DMSO. Mice that did not receive CAR T cells served as untreated controls (n = 4–6 mice per group).

(D–G) Mean (D and F) and individual (E and G) tumor growth curves following adoptive cell transfer (ACT; n = 4–5 per group). The dashed line indicates a tumor volume of 1,000 mm<sup>3</sup>, above which mice were evaluated for euthanasia according to the criteria described in Figure 4A. NR, no response; PR/CR, partial or complete response. Data represent different independent experiments using CAR T cells manufactured from distinct healthy donors. \**P* < 0.05, \*\**P* < 0.01, \*\*\**P* < 0.001, \*\*\*\**P* < 0.0001 using the log-rank (Mantel–Cox) test for survival analysis (A–C) and two-way repeated-measures ANOVA with Geisser–Greenhouse and Šídák correction (D and F). Error bars indicate the s.e.m.
